# Foxp3 depends on Ikaros for control of regulatory T cell gene expression and function

**DOI:** 10.1101/2023.05.23.541951

**Authors:** Rajan M. Thomas, Matthew C. Pahl, Liqing Wang, Struan F. A. Grant, Wayne W. Hancock, Andrew D. Wells

## Abstract

Ikaros is a transcriptional factor required for conventional T cell development, differentiation, and anergy. While the related factors Helios and Eos have defined roles in regulatory T cells (Treg), a role for Ikaros has not been established. To determine the function of Ikaros in the Treg lineage, we generated mice with Treg-specific deletion of the Ikaros gene (*Ikzf1*). We find that Ikaros cooperates with Foxp3 to establish a major portion of the Treg epigenome and transcriptome. Ikaros-deficient Treg exhibit Th1-like gene expression with abnormal expression of IL-2, IFNg, TNFa, and factors involved in Wnt and Notch signaling. While *Ikzf1*-Treg-cko mice do not develop spontaneous autoimmunity, Ikaros-deficient Treg are unable to control conventional T cell-mediated immune pathology in response to TCR and inflammatory stimuli in models of IBD and organ transplantation. These studies establish Ikaros as a core factor required in Treg for tolerance and the control of inflammatory immune responses.

## Introduction

The zinc finger DNA binding protein Ikaros is expressed in hematopoietic precursors, where it regulates genes involved in antigen receptor recombination, hemoglobin synthesis, and genome stability by recruiting chromatin remodeling complexes^1^. Germline deletion of *Ikzf1* in mice results in arrested erythroid and lymphoid development, leading to immunodeficiency and immature B and T cell leukemia. In conventional CD4+ and CD8+ T cells, Ikaros functions as a transcriptional repressor of inflammatory cytokine genes^2–5^. Conventional CD4+ T cells with loss of Ikaros function are unable to differentiate into peripherally-induced regulatory T cells (iTreg)^6^, and are resistant to suppression by thymic regulatory T cells^6^. However, the role of Ikaros in thymic Treg development and function has not been addressed.

The Ikaros family members Helios and Eos have each been deleted or knocked down in human or mouse regulatory T cells^7–9^. Loss of Eos function in Treg is associated with increased expression of inflammatory cytokines like IL-2 and IFNg, and an inability to control pathogenic conventional T helper cell responses in an IBD model^8,9^, although a separate study found that Eos-deficient Treg had normal function^10^. Helios also contributes to the control of Treg activation and cytokine production^11^, but this may be secondary to its role in promoting stable expression of the *Foxp3* gene^7,11,12^.

In this study, we conduct a genome-scale multi-omic analysis of open chromatin, active histone marks, Ikaros occupancy, Foxp3 occupancy, and gene expression in wild-type and *Ikzf1*-deficient regulatory T cells. We find that Ikaros plays a crucial role in establishing the normal landscape of enhancer activity, Foxp3 binding, and gene expression in Treg that cannot be filled by other Ikaros family members. Loss of Ikaros function in Treg results in uncontrolled T cell-dependent inflammatory responses *in vivo*.

## Results

To address the role of Ikaros in the Treg lineage, we crossed mice with a floxed *Ikzf1* allele^13^ with mice carrying a Foxp3-YFP-Cre reporter/driver. This strain generates an *Ikzf1*-null allele and the Cre neither generates a Foxp3 fusion protein nor affects Foxp3 function^14^. Male *Ikzf1*-fl-Foxp3-YFP-Cre mice (B6 background) do not develop overt autoimmune pathology under specific pathogen-free housing conditions over an 8 week timeframe, and basic aspects of T cell and Treg thymic development are unaltered (**Supplementary Figure 1a**). CD4+CD25+Foxp3+ peripheral Treg in these mice exhibit a nearly complete loss of Ikaros protein (**Figure 1a,b**) and full DNA demethylation at the *Foxp3* CNS2-TSDR (**Supplementary Figure 1b**), indicating that they are of thymic origin^15^. The expression of Ikaros in conventional CD4+ T cells from *Ikzf1*-fl-Foxp3-YFP-Cre mice is indistinguishable from that of control Foxp3-YFP-Cre mice (**Figure 1b**). *Ikzf1*-fl-Foxp3-YFP-Cre mice exhibit a statistically significant increase in total and effector Treg pools, with a concomitant decrease in the naive Treg pool in the spleen and lymph nodes (**Supplementary Figure 1c**). *Ikzf1*-deficient Treg express normal levels of Foxp3, GITR, and PD1 (**Figure 1c,d**), and higher levels of the high-affinity IL-2 receptor CD25 and the costimulatory receptor ICOS (**Figure 1c**). Ikaros-deficient Treg maintained Eos, Aiolos and Helios protein expression, exhibiting a mild increase in the expression of Eos and Aiolos particularly in the lymph nodes (**Figure 1d**).

**Figure 1.**
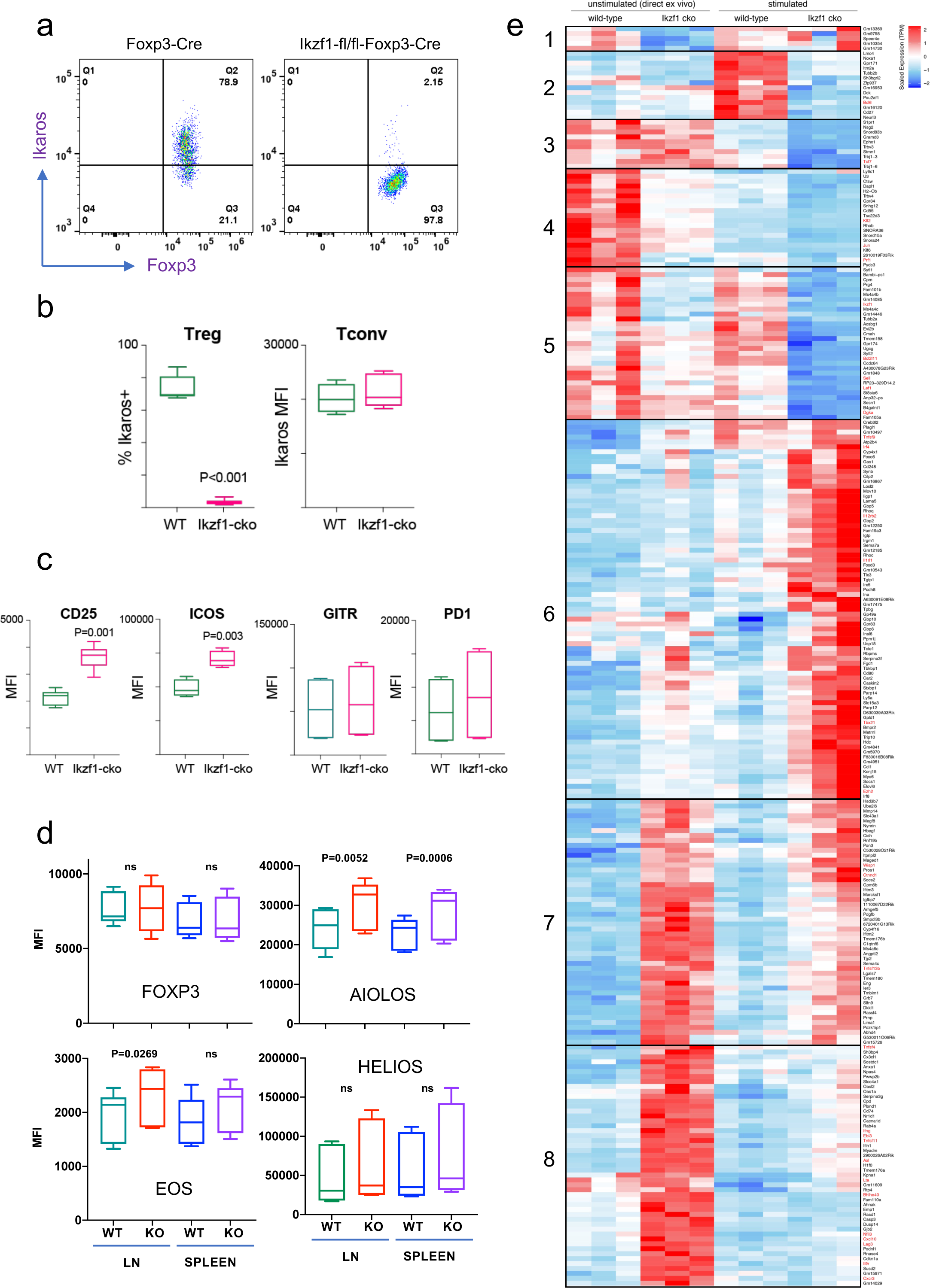
Impact of loss of Ikaros function on peripheral Treg phenotype. Example histograms (**a**) and quantified expression (**b**) of Ikaros protein by peripheral Treg and Tconv from Foxp3-YFP-Cre (green) and *Ikzf1*-fl-Foxp3-YFP-Cre (red) mice (N=6 mice per group). (**c**) Expression of CD25, ICOS, GITR, and PD1 by WT (green) and *Ikzf1*-cko (red) Treg (N=6 mice per group). (**d**) Flow cytometric measurement of Foxp3, Aiolos, Eos, and Helios protein expression in WT and *Ikzf1*-cko Treg (N=6 mice per group). (**e**) Transcriptomic analysis of WT vs. *Ikzf1*-cko Treg gene expression. Top differentially expressed genes (FDR<0.05) organized into 8 clusters in *ex vivo* or *in vitro* stimulated WT and *Ikzf1*-cko Treg. The heatmap represents scaled transcripts per million (tpm, N=3 replicates per group).

### Ikaros contributes significantly to the Treg gene expression program

Ikaros is a transcription factor, so to assess how loss of Ikaros function impacts the Treg gene expression program, we compared the transcriptomes of wild-type and Ikaros-deficient Treg isolated directly *ex vivo* from 6-8 week old aged-matched mice (N=3). A total of 661 genes were differentially expressed in *Ikzf1* cko Treg, 149 of which greater than 2-fold (FDR<0.05, **Supplementary Table 1**). Some of these genes were downregulated compared to wild-type Treg (**Figure 1e**, clusters 1, 4 and 5), but 80% were upregulated (**Figure 1e**, clusters 7 and 8), indicating that Ikaros functions primarily as a transcriptional repressor in Treg. Loss of Ikaros results in up-regulation of at least 12 factors that negatively regulate Treg function, *e.g*., multiple genes involved in Wnt signaling (*Wisp1, Ctnnd1, Ctnna1*), *Ox40, Tlr2, Lag3, Tnf,* and *Ifng* (**Figure 2a** and **Supplementary Table 2**). Increased *Ifng* expression correlated with decreased DNA methylation at the *Ifng* locus compared to wild-type Treg (**Supplementary Figure 1d**). *Ikzf1* cko Treg also exhibited down-regulation of at least 10 factors that are required for full Treg function, *e.g*., the activin receptor *Acvr1b, Nr4a1/Nur77, Tet1*, and perforin (**Figure 2b** and **Supplementary Table 2**). *Bcl6*, which is required for follicular Treg function^16^, is also down-regulated in Ikaros-deficient Treg. However, *Ikzf1* cko Treg up-regulated at least 24 factors known to promote Treg function (**Supplementary Table 2**), suggesting that the loss-of-function program may be counteracted by a gain-of-function program in the absence of inflammation.

**Figure 2.**
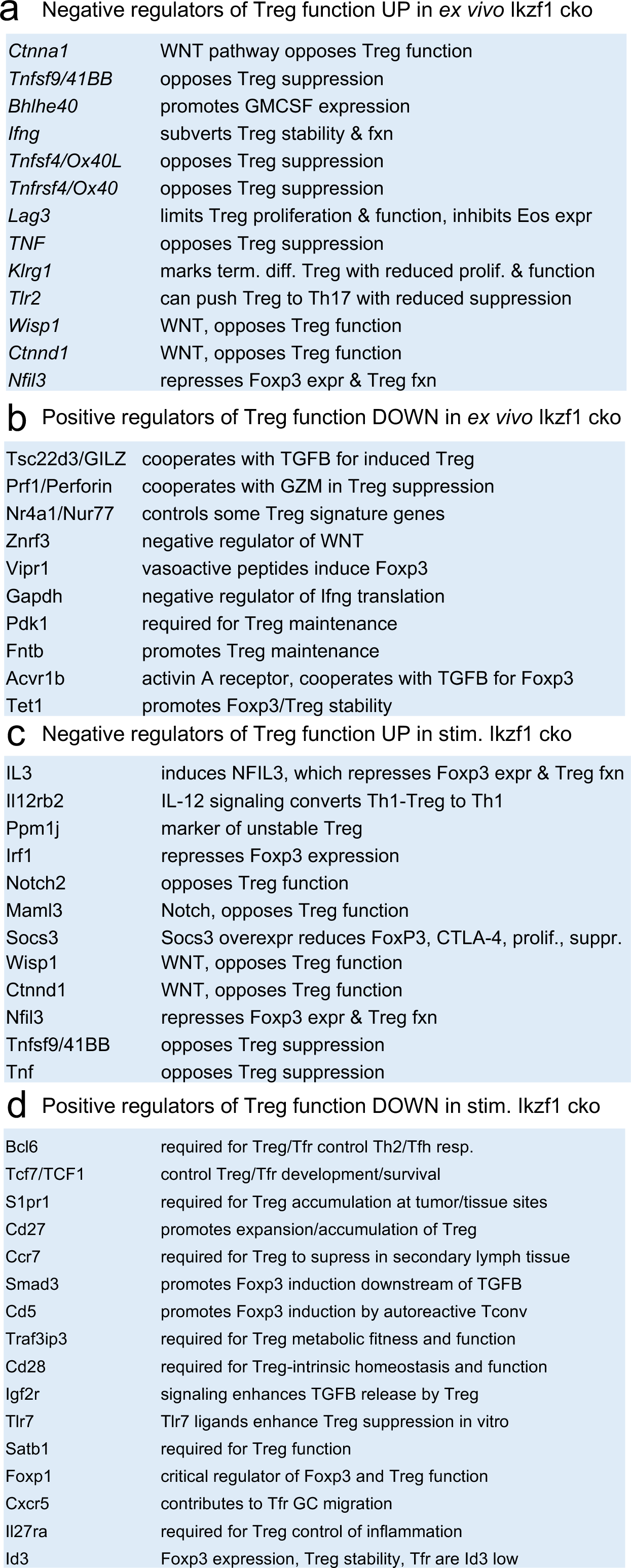
Survey of relevant differentially expressed genes in WT vs. *Ikzf1*-cko Treg. (**a**) Known negative regulators of Treg function up-regulated in *ex vivo Ikzf1*-cko Treg, (**b**) positive regulators of Treg function down-regulated in *ex vivo Ikzf1*-cko Treg, (**c**) negative regulators of Treg function up-regulated in *in vitro stimulated Ikzf1*-cko Treg, (**d**) positive regulators of Treg function down-regulated in *in vitro stimulated Ikzf1*-cko Treg. See **Supplementary Table 1** for all differential genes and **Supplementary Table 2** for a larger list of functionally relevant differentially expressed genes.

To simulate antigen encounter by Treg during an immune response, we stimulated wild-type and Ikaros-deficient Treg through the TCR and CD28 *in vitro*. Previous studies established a role for Ikaros in restricting IL-2 and IFNg production by conventional CD4+ and CD8+ T cells^2–5^. We find that Ikaros plays a similar role in Treg, as unlike wild-type Treg, *Ikzf1* cko Treg are capable of secreting IL-2 and IFNg protein upon stimulation (**Figure 3a**). Treg from mice expressing a dominant-negative form of Ikaros likewise ectopically express IL-2, IFNg, and TNFa at the protein level upon stimulation (**Figure 3b**). Consistent with their increased expression of IL-2 and IL-2R, Ikaros-deficient Treg exhibit enhanced IL-2-induced STAT5 phosphorylation (**Figure 3c**) and increased proliferative capacity (**Figure 3d**) compared to wild-type Treg.

**Figure 3.**
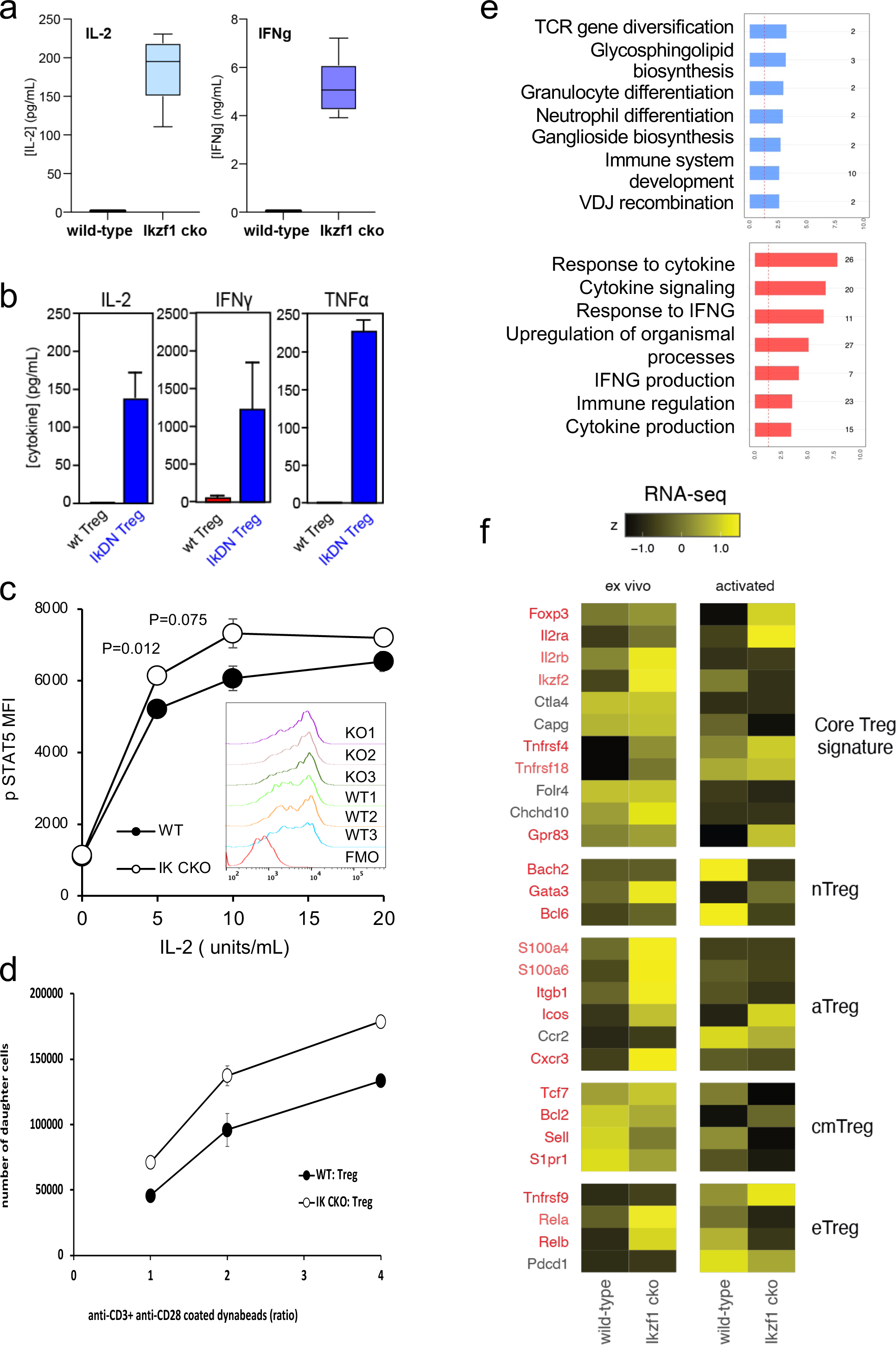
*In vitro* analysis of WT vs. *Ikzf1*-cko Treg. (**a**) Secretion of IL-2 and IFNg protein by WT and *Ikzf1*-cko Treg measured by ELISA (N=3). (**b**) IL-2, IFNg, and TNFa production by Treg from WT mice vs. mice with a dominant-negative form of Ikaros (IkDN) measured by ELISA (N=3). (c) IL-2-induced phosphorylation of STAT5 in WT and *Ikzf1*-cko Treg measured by flow cytometry *in vitro*. Mean fluorescence intensity (MFI) and individual histograms (inset, N=3) are depicted. (d) Activation-induced proliferation of WT (closed) and *Ikzf1*-cko (open) Treg measured by dye dilution (N=3). (**e**) Gene ontology analysis of genes down-regulated (top panel) and up-regulated (bottom panel) in *Ikzf1*-cko compared to WT Treg. The x-axis is fold enrichment and numbers to the right are unique genes in each pathway. (**f**) Differential expression of core Treg, nTreg, aTreg, cTreg, and eTreg genes^19^ in *ex vivo* (left panel) and *in vitro* stimulated (right panel) WT and *Ikzf1*-cko Treg. The heatmap represents z-score and genes significantly differentially expressed are shown in red.

At genome scale, stimulation through the TCR and CD28 led to differential expression of 895 genes (FDR<0.05, **Supplementary Table 1**), 533 down-regulated (**Figure 1e**, clusters 2, 3 and 5) and 362 up-regulated in *Ikzf1* cko Treg compared to wild-type Treg (**Figure 1e**, clusters 6, 7 and 8). Consistent with its role in antigen receptor selection during lymphocyte development^17^, gene ontology analysis (**Supplementary Table 3**) of genes down-regulated in Ikaros-deficient Treg shows enrichment for immune cell development and TCR/VDJ recombination and diversification (**Figure 3e**). Genes upregulated in Ikaros-deficient Treg are enriched for networks involved in interferon and cytokine production and responses (**Figure 3e**). The set of upregulated genes includes at least 20 factors known to promote Treg function like *Foxp3, Il2ra, Icos, Ezh2,* and *Gpr83* (**Supplementary Table 2**). However, loss of Ikaros results in up-regulation of at least 11 factors that negatively regulate Treg function like *Irf1, Il12rb2* (IL-12 receptor), *Il3*, and several genes in the Notch and Wnt pathways like *Notch2, Maml3, Rbpj, Wisp,* and *Ctnnd1* (**Figure 2c**), and down-regulation of at least 14 factors that are required for full Treg function like *Tcf7, Satb1, Foxp1, Id3, Smad3, Il27ra, Tlr7*, and follicular Treg (Tfreg) genes like *Bcl6, Cxcr5,* and *S1pr1* (**Figure 2d**).

The Wnt pathway is normally repressed in Treg, and ectopic Wnt signaling in Treg has been associated with ectopic IFNg production and reduced suppressive function^18^. We show that the Wnt signaling factor B-catenin is elevated in *Ikzf1* cko Treg (**Figure 4a**), and that ectopic IFNg production by Ikaros-deficient Treg is Wnt-dependent, while IFNg production by conventional CD4+ T cells does not depend on Wnt (**Figure 4b,c**). Together, these transcriptomic analyses indicate that Ikaros normally regulates a large proportion of the core Treg program^19^ (**Figure 3f**, genes in red), and is required to restrain Wnt, Notch, and inflammatory cytokine gene expression in the Treg lineage. The impact of the loss of Ikaros function on the Treg transcriptome likely stem from both direct, cell-intrinsic effects and from indirect, cell-extrinsic effects.

**Figure 4.**
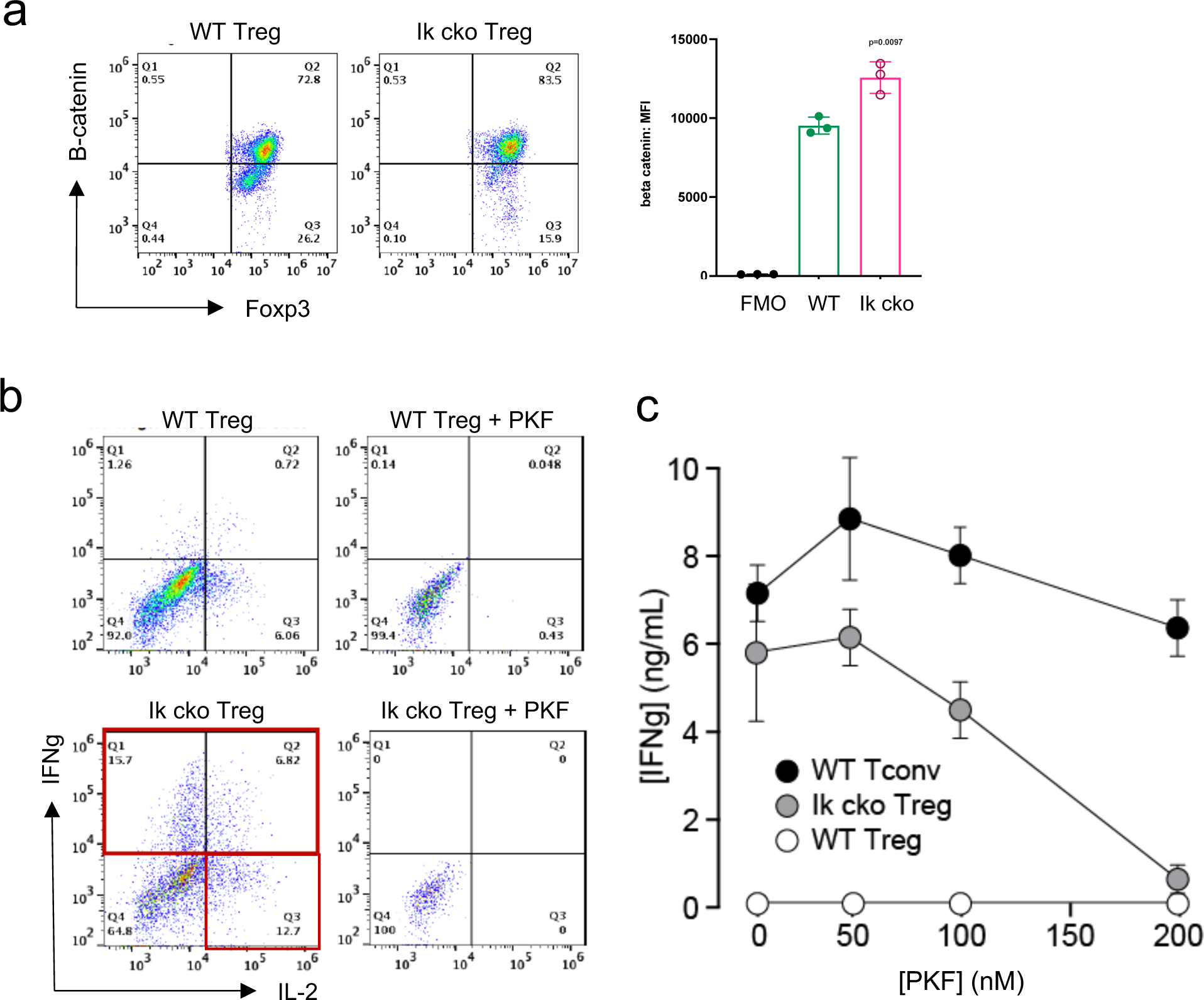
Ectopic activation of the Wnt-catenin pathway in *Ikzf1*-cko Treg. (**a**) Expression of B-catenin by WT (left histogram) vs. *Ikzf1*-cko (right histogram) Treg measured by flow cytometry (plot depicts B-catenin MFI from N=3 experiments). (**b**) IFNg secretion by WT (top panels) vs. *Ikzf1*-cko (bottom panels) Treg activated with (right panels) or without (left panels) the Wnt pathway inhibitor PKF. (**c**) IFNg secretion (measured by ELISA) by Ikzf1-cko Treg, but not by conventional T cells, is inhibited in a dose-dependent manner by PKF (N=3).

### Ikaros is required for establishing the Treg open chromatin and enhancer landscape

To gain mechanistic insight into how Ikaros regulates the Treg gene expression program, we measured Ikaros binding, Foxp3 binding, chromatin accessibility, and H3K27ac enhancer marks in wild-type and Ikaros-deficient Treg using ATAC-seq and ChIP-seq. Loss of Ikaros function induced remodeling of 1431 genomic regions (N=3, FDR<0.05, **Supplementary Table 4** and **Supplementary Figure 2a-c**), one-third of which (513) exhibiting reduced accessibility and two-thirds (918) of which become more accessible (**Figure 5a**). Regions with reduced accessibility in Ikaros-deficient Treg were enriched for nearby genes involved in leukocyte development and differentiation, while regions showing increased accessibility were enriched for genes involved in cytokine signaling and response to interferon-gamma (**Supplementary Figure 2d**). At genome scale, increased accessibility at genomic elements after deletion of Ikaros correlated significantly with increased expression of nearby genes, while decreased accessibility correlated significantly with reduced gene expression (**Supplementary Figure 2e**).

**Figure 5.**
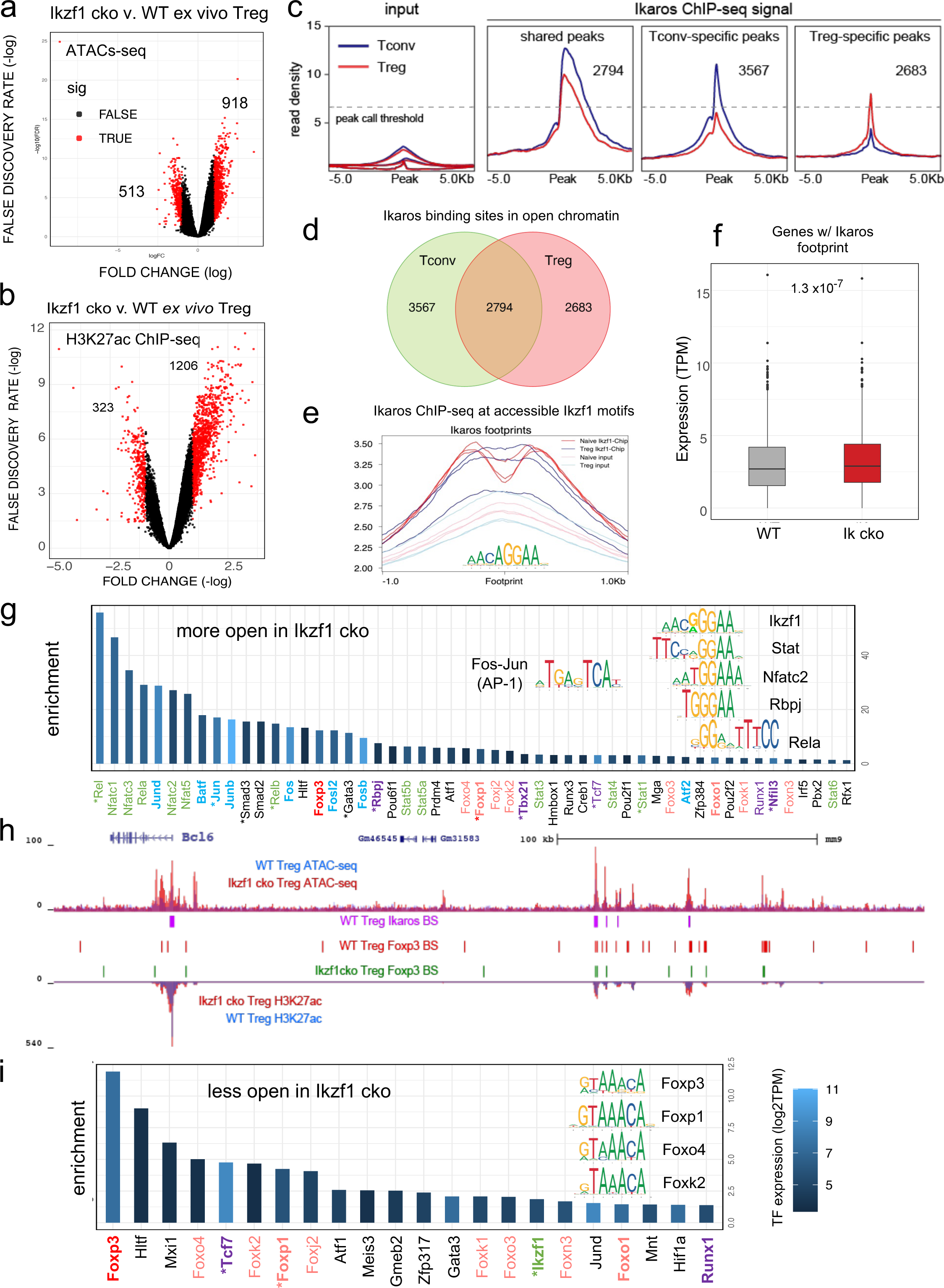
Genome-wide assessment of open chromatin, histone acetylation, and Ikaros occupancy in WT and *Ikzf1*-cko Treg. Differential analysis of open chromatin (**a**) and H3K27ac (**b**) in WT vs. *Ikzf1*-cko Treg (FDR<0.05, N=3). Peaks with FC>2 are depicted in red. (**c**) Ikaros ChIP-seq signal (read density) at genomic regions shared (panel 2) in Tconv (blue) vs. Treg (red), unique to Tconv (panel 3), or unique to Treg (panel 4). Panel 1 depicts read densities at the same regions in control input libraries. (**d**) Unique vs. shared Ikaros binding sites in Treg (red) vs. Tconv (green) open chromatin. (**e**) Enrichment of Ikaros ChIP-seq signal (footprint, solid lines) or input background (transparent lines) at accessible Ikaros motifs (AGGAA) in WT Treg (blue) and Tconv (red). (**f**) Expression (tpm) of genes with open chromatin enriched for the Ikaros consensus binding motif in WT vs. *Ikzf1*-cko Treg. (**g**) Enrichment of TF consensus binding motifs in genomic regions that are more accessible in *Ikzf1*-cko Treg. Inset depicts motifs for Ikaros/Ikzf1, Stat, Nfat, Rbpj, Rela, and AP-1. (**h**) Open chromatin (top tracks) and H3K27ac (bottom tracks) in WT (blue) and *Ikzf1*-cko (red) Treg, Ikaros binding sites (purple marks) and Foxp3 binding sites (red marks) in WT Treg, and Foxp3 binding sites in *Ikzf1*-cko Treg (green marks) at the *Bcl6* locus. (**i**) Enrichment of TF consensus binding motifs in genomic regions that are less accessible in *Ikzf1*-cko Treg. Inset depicts motifs for forkhead family members. In (**g**) and (**i**), factors with roles in Treg function are colored green and purple, forkhead family members are colored red, and factors differentially expressed in *Ikzf1*-cko Treg are indicated with an asterisk. ATAC-seq was performed on Treg purified directly *ex vivo*, while ChIP-seq analyses were performed on Treg expanded *in vivo* using IL-2/anti-IL-2 complexes.

To explore how Ikaros regulates the Treg enhancer landscape, we measured histone acetylation at nucleosomes flanking open chromatin regions in wild-type and *Ikzf1* cko Treg (N=3, **Supplementary Figure 2f,g**). Out of approximately 21,000 H3K27ac peaks called in both cell populations, 40% were affected by loss of Ikaros function (**Supplementary Table 5**), with 323 regions showing a >2-fold reduction acetylation and 1206 regions exhibiting a >2-fold increase in H3K27ac (**Figure 5b**). Differential analysis also showed that Ikaros controls histone acetylation at ∼25% of Treg open chromatin regions (OCR, 9937 out of ∼40,000, FDR <0.05), with the vast majority (78%, 7788) showing increased acetylation upon loss of Ikaros function (**Supplementary Table 5** and **Supplementary Figure 2h**). At genome scale, enrichment of the H3K27ac mark correlates with regional accessibility (**Supplementary Figure 2i**), and strength of the enhancer signature at a given element correlates with the level of nearby gene expression (**Supplementary Figure 2j**). Dense collections of multiple enhancers that tend to drive expression of genes involved in cell identity are called ‘super-enhancers’. We defined 1,700 Treg super-enhancers based on H3K27ac density (**Supplementary Table 6**), 20% of which (324) are regulated by Ikaros (**Supplementary Figure 2k**).

Ikaros ChIP-seq analysis identified 7642 Ikaros binding sites in WT Tconv, 83% of which (6361) are located in open chromatin, and 7061 Ikaros binding sites in WT Treg, 76% of which (5477) are located in open chromatin (**Figure 5c** and **Supplementary Table 7**). Of all accessible Ikaros binding sites detected, 39% (3567) were Tconv-specific, 31% (2794) were shared between Tconv and Treg, and 29% (2683) were only detected in Treg (**Figure 5d**). The Ikaros ChIP-seq signal is enriched at accessible Ikaros motifs in the Treg and Tconv genome (**Figure 5e**), and the set of genes with accessible Ikaros binding motifs showed increased expression in Ikaros-deficient compared to wild-type Treg (**Figure 5f**). Motif analysis (**Supplementary Table 8**) at regions that exhibit increased accessibility in *Ikzf1* cko Treg shows enrichment of the Ikaros GGGAA core binding sequence that is shared with immune trans-activators like NFkB, NFAT, Notch, and Stat1/4, and these regions are also enriched for AP-1 (Fos/Jun) and T-bet (Tbx21) motifs (**Figure 5g** inset). This suggests that Ikaros can directly repress inflammatory gene expression in Treg by competing with NFkB, NFAT, Notch, and Stats for binding to enhancers and recruiting epigenetic factors that silence these elements^20–24^. Loci under direct repressive control of Ikaros in Treg include *Bcl6*, *Notch2, Irf4,* and *Ifng* (**Figure 5h** and **Supplementary Figure 3a-c**). However, the majority of genomic regions exhibiting increased accessibility in *Ikzf1* cko Treg are not bound by Ikaros in wild-type cells (873 of 918, **Supplementary Figure 3d**), suggesting that indirect gene regulation due to observed alterations in the expression of other transcription factors is another mechanism by which Ikaros establishes the Treg gene expression program.

### Foxp3 cooperates with Ikaros for DNA binding across the Treg genome

Regions that exhibit reduced accessibility in *Ikzf1* cko compared to wild-type Treg are enriched for the binding motif for Ikaros and the motif for TCF1 (**Figure 5g**), a factor that cooperates with Foxp3 to enforce Treg function^25^ and is down-regulated in *Ikzf1* cko Treg (**Figure 3f**). These regions are likewise enriched for the GTAAACA Foxp3/forkhead motif (**Figure 5i** inset), suggesting that Foxp3 may cooperate with Ikaros at these sites. To test this, we compared Foxp3 genome occupancy in wild-type vs. *Ikzf1* cko Treg by ChIP-seq (**Figure 6a**, **Supplementary Figure 4a-c**). A total of 4423 Foxp3 binding sites were detected in the open chromatin landscape of wild-type Treg (**Supplementary Table 9**), and this ChIP-seq signal was enriched at accessible Foxp3 motifs. Consistent with the motif analyses (**Figure 5h**), we find a remarkable 74% of all Foxp3 binding sites are co-bound by Ikaros (3255 of 4423 sites, **Figure 6b**). Loss of Ikaros in *Ikzf1* cko Treg results in reduced Foxp3 binding affinity at 70% of Ikaros-Foxp3 co-bound sites (2254 of 3256, **Figure 6b** inset), and reduced Foxp3 binding at 80% of all sites strongly bound by Foxp3 in wild-type Treg (3543 of 4422, **Figure 6a,c**). As a result, the set of all direct target genes with accessible Foxp3 binding motifs showed significantly increased expression in Ikaros-deficient compared to wild-type Treg (**Figure 6d**).

**Figure 6.**
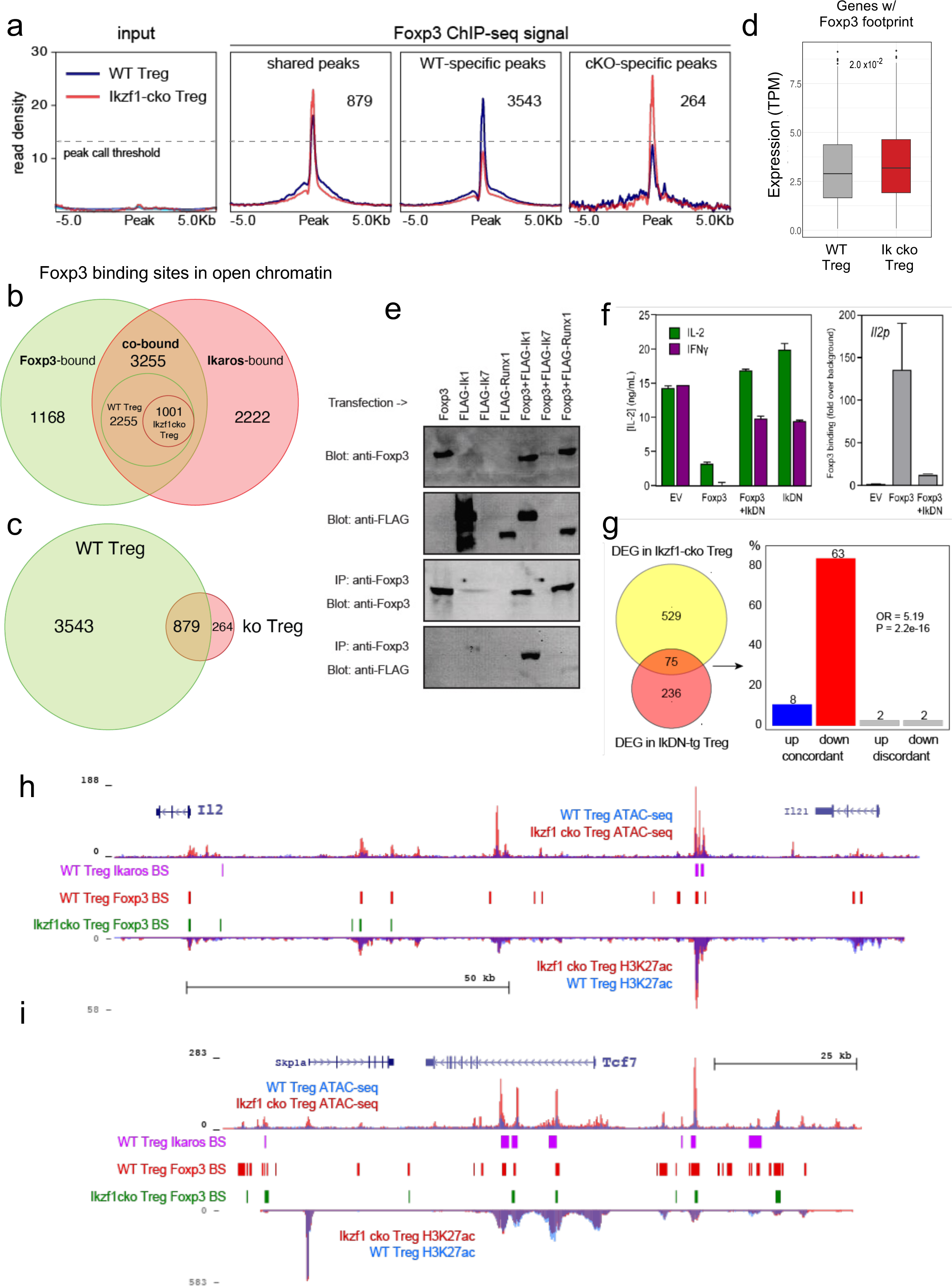
Ikaros-dependent Foxp3 function in Treg. (**a**) Input vs. Foxp3 ChIP-seq at genomic regions shared or unique in WT (blue) vs. *Ikzf1*-cko (red) Treg (N=3 per group). (**b**) Foxp3- (green), Ikaros- (red) and Foxp3-Ikaros co-bound (orange) OCR. Inset depicts Foxp3-Ikaros co-bound regions in WT (green) vs. *Ikzf1*-cko (red) Treg. (**c**) Foxp3 binding sites in WT (green) vs. *Ikzf1*-cko (red) Treg. (**d**) Expression (tpm) of genes enriched for accessible Foxp3 consensus motifs in WT vs. *Ikzf1*-cko Treg. (**e**) 293T cells transfected with FLAG-tagged full-length Ikaros (Ik1), DNA-binding mutant Ikaros (Ik7), or Runx1 alone (lanes 1-4) or in combination with untagged Foxp3 (lanes 5-7). Whole extracts (panels 1-2) or Foxp3-immunoprecipitated extracts (panels 3-4) probed for Foxp3 or FLAG. (**f**) IL-2 or IFNg production (left panel) and Foxp3 ChIP-qPCR at *Il2* promoter (right panel) in Tconv transduced with vector, Foxp3, Ik7/DN, or Foxp3+Ik7/DN. (**g**) Concordant vs. discordant genes co-regulated in *Ikzf1*-cko vs. IkDN Treg (odds ratio=5.19, P value=2.2×10^-16^). (**h,i**) Open chromatin (top) and H3K27ac (bottom) in WT (blue) and *Ikzf1*-cko (red) Treg, Ikaros binding (purple marks) and Foxp3 binding (red marks) in WT Treg, and Foxp3 binding in *Ikzf1*-cko Treg (green marks) at *Il2* (**h**) and *Tcf7* (**i**).

Consistent with these observations, we find that Ikaros and Foxp3 exist in a complex in the nuclei of cells (**Figure 6e**), confirming prior proteomic data suggesting a physical interaction between Foxp3 and Ikaros in transfected cells^26^. At the Foxp3 and Ikaros co-bound target genes *Il2* and *Ifng*, retroviral expression of Foxp3 in CD4+ T cells results in direct promoter occupancy and silencing of both genes (**Figure 6f**). Co-expression of dominant-negative Ikaros (IkDN or Ik7) abrogates binding of Foxp3 to the *Il2* promoter and inhibits the repressive activity of Foxp3 (**Figure 6f**). At genome scale, a significant number of genes in addition to *Il2* are concordantly regulated by both dominant-negative Ikaros and *Ikzf1* gene deletion (**Figure 6g**). In addition to the promoter, we also observe Ikaros-Foxp3 co-binding at a defined distal enhancer of *Il2* located 83 kb upstream of the promoter^27^ in wild-type Treg (**Figure 6h**). Loss of Ikaros function in *Ikzf1* cko Treg results in loss of Foxp3 binding at this enhancer, which is accompanied by increased histone acetylation, chromatin accessibility (**Figure 6h**), and *Il2* expression (**Figures 1** and **2**). Other examples of Foxp3-regulated genes that exhibit Ikaros-dependent Foxp3 binding, enhancer activity and expression are *Tcf7* (**Figure 6i**), *Il2ra, Rbpj,* and *Maml3* (**Supplementary Figure 4d,e**). Together, these results indicate that a large portion of the Treg epigenome and transcriptome, including two-thirds of the core Treg program and the majority of the Foxp3 gene regulatory program, is dependent on Ikaros.

### Ikaros is required for Treg control of conventional T cell differentiation

The large-scale dysregulation of gene expression in *Ikzf1*-cko Treg, especially upon stimulation, suggests that extrinsic control of inflammatory immune responses may be dysregulated in mice lacking Ikaros in the Treg lineage. We observed no clear signs of frank autoimmunity in aged (1 year-old) *Ikzf1*-fl-Foxp3-YFP-Cre mice (**Supplementary Figure 5a**). The conventional T cell pool in 6-8 week-old (**Figure 7**) and 10 month-old (**Supplementary Figure 5**) *Ikzf1*-fl-Foxp3-YFP-Cre mice showed a statistically significant accumulation of CD4+ T cells (**Figure 7a, Supplementary Figure 5b**) with a memory phenotype (**Figure 7b,c**), and a concomitant reduction in naive phenotype CD4+ T cells (**Figure 7d, Supplementary Figure 5b**). Ikaros-deficient Treg maintained comparable suppressive activity against wild-type Tconv *in vitro* (**Figure 7e**), however, Ikaros-sufficient, conventional CD4+ T cells from mice lacking Ikaros in the Treg lineage were resistant to suppression by both *Ikzf1*-cko and wild-type Treg (**Figure 7f,g**), likely due increased frequency of suppression-resistant memory cells^28,29^. At 6 months, *Ikzf1*-fl-Foxp3-YFP-Cre mice also exhibited increased frequencies of follicular helper CD4+ T cells in the lymph nodes and spleen (**Figure 7h**), which was associated with elevated levels of total IgM, IgG, and especially IgA in the serum (**Figure 7i**). The elevated IgA was accompanied by increased frequencies of IgA-positive B cells in the spleen and mesenteric lymph nodes (**Figure 7j,k**). These results suggest perturbed immune homeostasis in mice that lack Ikaros function in Treg owing to a defect in the control of conventional CD4+ T cell differentiation.

**Figure 7.**
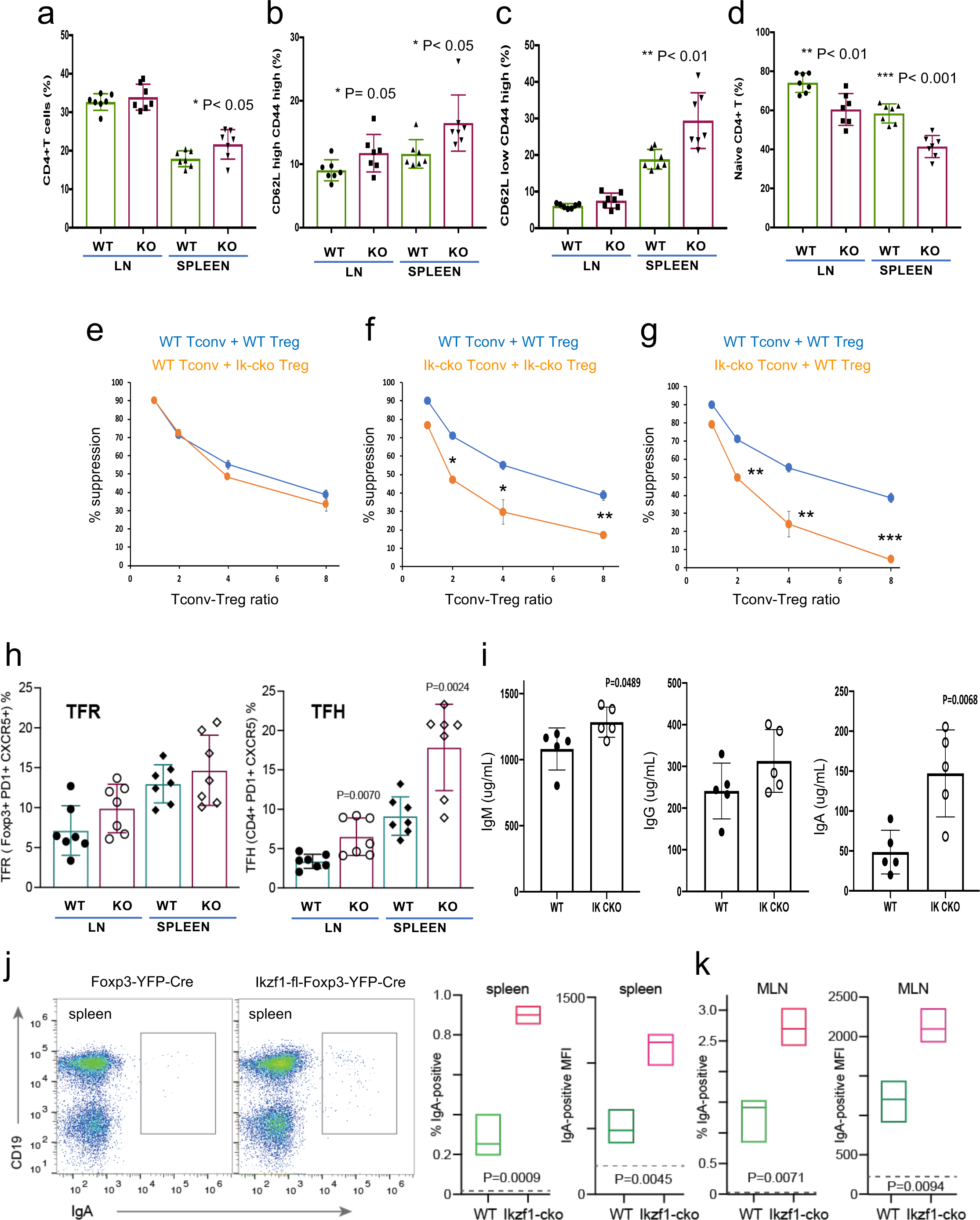
Immunophenotyping of Tconv from *Ikzf1*-fl-Foxp3-YFP-Cre and Foxp3-YFP-Cre mice. Frequencies of total (**a**), memory (**b,c**), and naïve (**d**) phenotype Tconv in secondary lymphoid tissues of 6-8 week-old *Ikzf1*-fl-Foxp3-YFP-Cre (purple) and Foxp3-YFP-Cre (green) mice (N=7). (**e**) *In vitro* suppressive activity of WT (blue) vs. *Ikzf1*-deficient (orange) Treg against Tconv from WT mice. (**f**) *In vitro* suppressive activity of WT Treg against Tconv from WT mice (blue) vs. *Ikzf1*-deficient Treg against Tconv from *Ikzf1*-cko mice (orange). (**g**) *In vitro* suppressive activity of WT Treg against Tconv from WT (blue) vs. *Ikzf1*-deficient mice (orange). Tconv proliferation was measured by dye dilution in all cultures (N=3). (**h**) Frequencies of Foxp3+PD1^hi^CXCR5^hi^ follicular regulatory T cells (Tfr) and PD1^hi^CXCR5^hi^ follicular helper T cells (Tfh) in secondary lymphoid tissues of 6 month-old *Ikzf1*-fl-Foxp3-YFP-Cre (purple) and Foxp3-YFP-Cre (green) mice (N=7). (**i**) Total serum levels of IgM, IgG, and IgA from *Ikzf1*-fl-Foxp3-YFP-Cre vs. Foxp3-YFP-Cre mice (N=4). Frequency of IgA-positive B cells and surface density (MFI) of IgA-positive B cells in spleen (**j**) and mesenteric lymph nodes (**k**) from 9 month-old WT (green) vs. *Ikzf1*-deficient (red) mice (N=3). P values are indicated for significant differences.

### Ikaros is required for Treg control of pathogenic T cell-mediated mucosal inflammation and acquired immune tolerance

To address this, we tested the ability of Ikaros-deficient Treg to control inflammatory colitis in an *in vivo* adoptive transfer model of IBD. Rag-deficient mice that received conventional CD4+CD25-negative T cells alone (N=5) developed severe disease as evidenced by progressive weight loss (**Figure 8a**), gross and histological intestinal pathology (**Figure 8b** and **Supplementary Figure 6a**), extensive cellular infiltration and tissue damage in the inner mucosal epithelial layer of the colon (**Figure 8c**), and high numbers of activated Tconv in the colon (**Figure 8c**), spleen and mesenteric lymph nodes of these animals (**Supplementary Figure 6d-g**). Upon co-transfer into Rag-deficient mice (N=5), wild-type Treg accumulated in the mesenteric lymph nodes (**Supplementary Figure 6i**) and colon (**Figure 8d**), were able to control the activation and expansion of conventional helper T cells in lymphoid tissues (**Supplementary Figure 6e,g**), and limit CD4+ T cell infiltration into the intestinal epithelium (**Figure 8d**). Recipients of wild-type Treg exhibited only mild intestinal pathology at the gross and microscopic levels (**Figure 8b,d** and **Supplementary Figure 6b**), and lost little weight over the course of the experiment (**Figure 8a**). Co-transferred Ikaros-deficient Treg accumulated in the spleen (**Supplementary Figure 6h,i**) and, despite reduced expression of the alpha4 beta7 integrin (*Itga4, Itgb7*) and *Ccr9* genes involved in homing to the intestine, accumulated in the intestinal epithelium to numbers 5-10-fold higher than in the recipients of wild-type Treg (**Figure 8e**). Despite their presence of large numbers in the intestinal mucosa, Ikaros-deficient Treg were completely unable to protect RAG-deficient mice (N=5) from infiltration and colitis mediated by wild-type conventional T cells at the level of weight loss (**Figure 8a**) and intestinal pathology (**Figure 8b,e** and **Supplementary Figure 6c**). Similar to mice that received no Treg, recipients of Ikaros-deficient Treg exhibited extensive inflammatory infiltrates in the colon, with thickening and detachment of the epithelial layer from underlying tissues (**Figure 8e**). Cell necrosis and mononuclear lymphocyte infiltration were also more pronounced in recipients of Ikaros-deficient Treg. These results indicate that Treg depend on Ikaros to control mucosal inflammation during a conventional T cell response.

**Figure 8.**
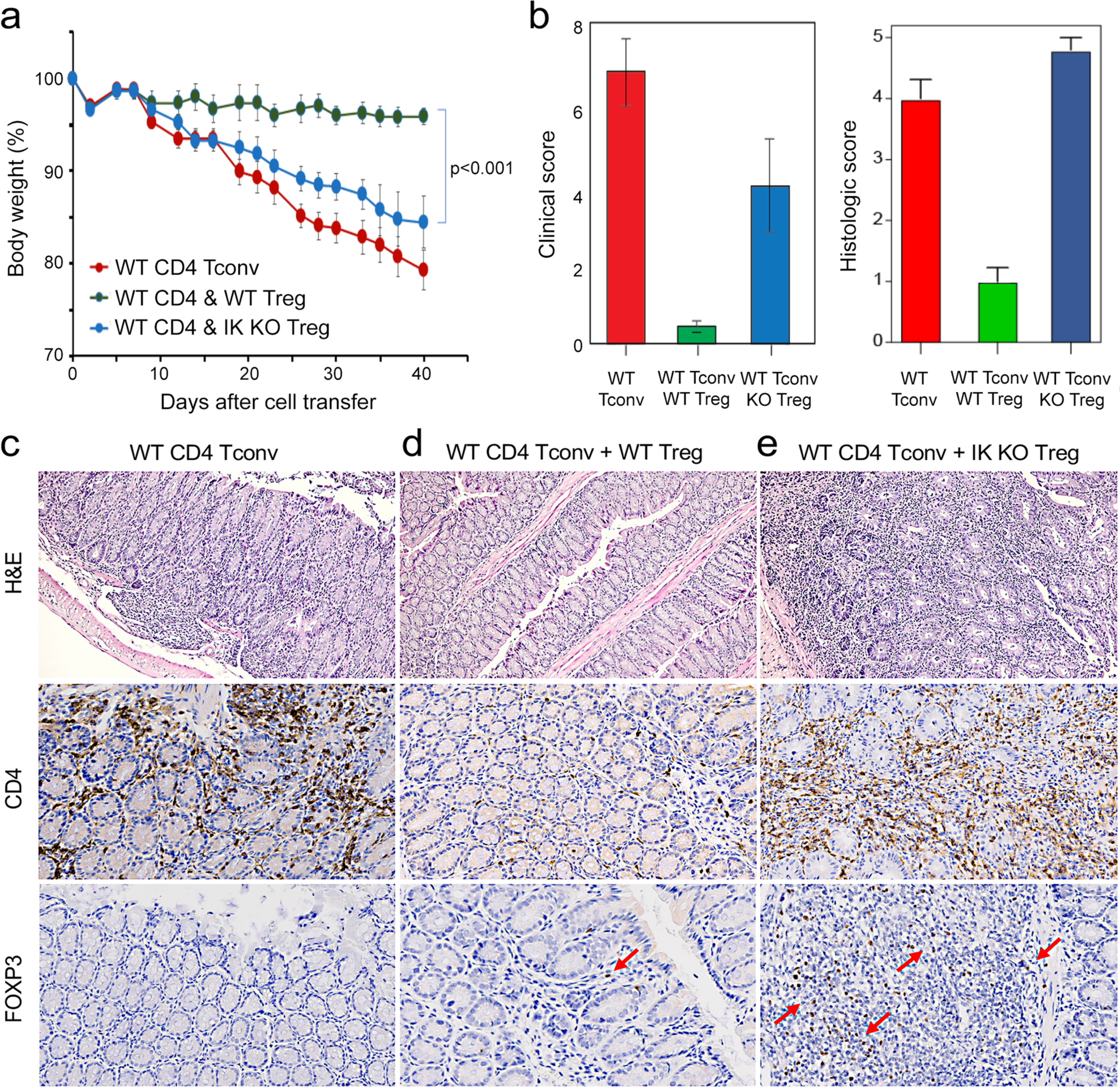
Role of Ikaros in Treg-mediated control of inflammatory colitis. WT CD4+CD25-Tconv were transferred alone (red), or together with WT (green) or *Ikzf1*-cko (blue) CD4+CD25+ Treg into RAG1ko mice (N=5). Animal weight was monitored for 40 days (**a**), and intestines were scored for pathology at the gross and histologic levels (**b**). Example histopathology of colons from RAG1ko recipients of WT Tconv (**c**), WT Tconv + WT Treg (**d**), and WT Tconv + *Ikzf1*-cko Treg (**e**). H&E (top row), CD4 (middle row), and Foxp3 (bottom row) staining are shown at 200x. Mean Foxp3+ cells per 200X field from N=3 animals is 3.4 in (d) and 22 in (e), P<0.05.

Regulatory T cells are required for the induction of peripheral alloimmune tolerance. To determine whether Treg-intrinsic Ikaros function is required for acquired tolerance to organ transplants, we transplanted fully mismatched cardiac allografts into *Ikzf1*-fl-Foxp3-YFP-Cre or control Foxp3-YFP-Cre mice under combined blockade of the CD28 and CD40 costimulatory pathways (N=5 per group). While costimulatory blockade induced long-term allograft tolerance in wild-type recipients, this treatment failed to induce tolerance in mice lacking Ikaros in the Treg lineage (**Figure 9a**). Similar results were obtained when anti-CD40L plus donor-specific transfusion was used as a tolerizing regimen (**Supplementary Figure 6j,k**). Intragraft analysis of gene expression showed elevated levels of multiple Th1-related transcripts in rejecting grafts from *Ikzf1*-Treg-cko recipients, despite elevation of Foxp3 (**Figure 9b**). Histopathological analysis of cardiac graft tissue from *Ikzf1*-Treg-cko recipients showed extensive myocardial necrosis (**Figure 9c**) associated with increased CD4+ T cell infiltration despite numbers of Foxp3+ Treg comparable to that in tolerant recipients (**Figure 9d**). These results reveal an important role for Ikaros in the ability of Treg to control inflammation and establish acquired immune tolerance.

**Figure 9.**
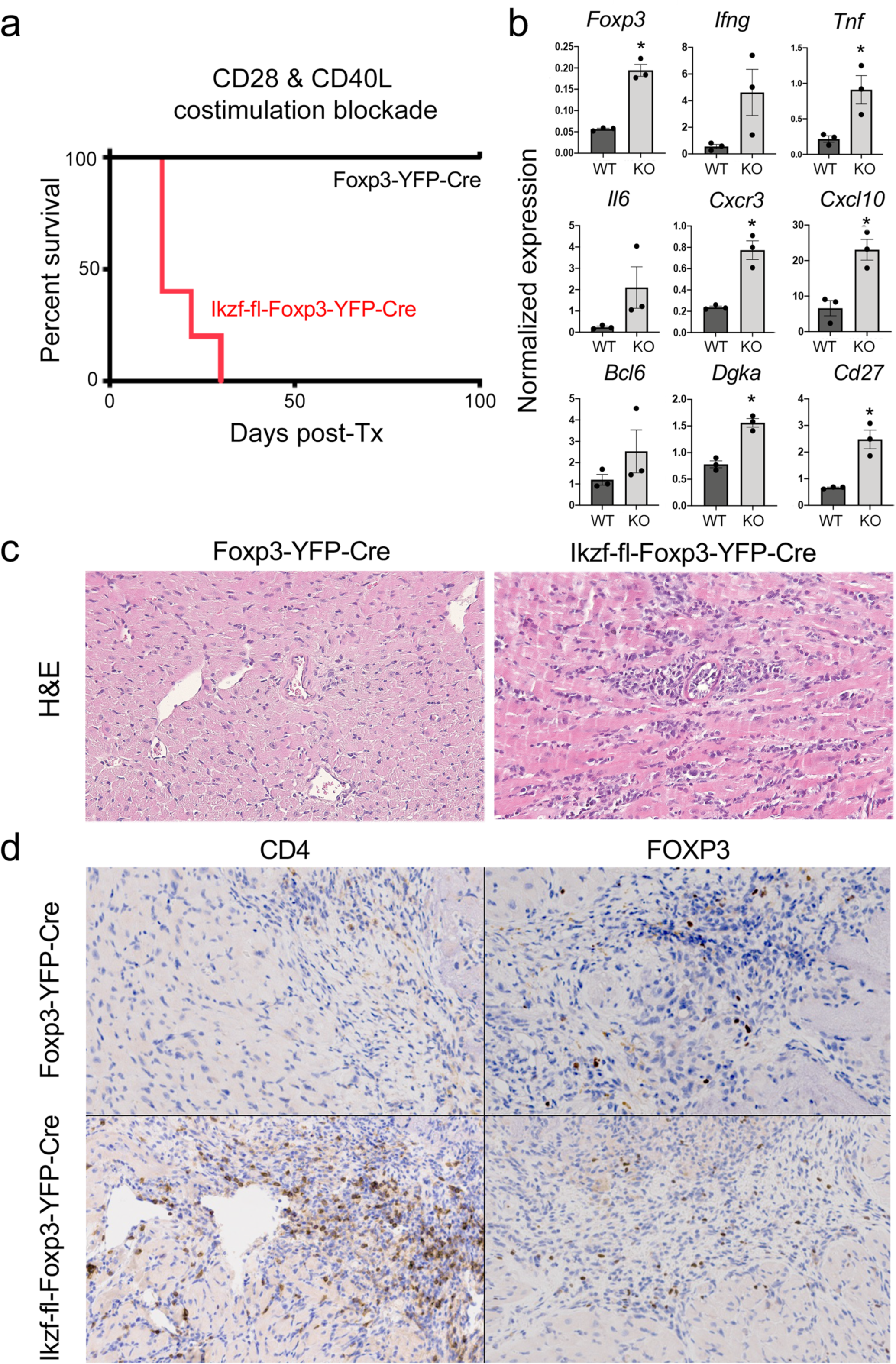
Role of Ikaros in Treg-dependent acquired cardiac transplant tolerance. (**a**) B6 *Ikzf1*-fl-Foxp3-YFP-Cre (red) or Foxp3-YFP-Cre (black) mice (N=5) received BALB/c cardiac allografts under combined CD28+CD40 costimulatory blockade and graft survival was monitored for 100 days. (**b**) Analysis of intra-graft transcript levels of the indicated genes from grafts harvested at day 19 post-transplant. Histopathological analysis of cardiac grafts harvested at day 19 post-transplant from Foxp3-YFP-Cre and *Ikzf1*-fl-Foxp3-YFP-Cre recipients by H&E (**c**) and CD4 and Foxp3 (**d**) staining (scale=200x).

## Discussion

The studies reported here establish a crucial role for Ikaros in regulatory T cells that cannot be replaced by the Ikaros family members Helios, Eos and Aiolos. While Ikaros is required for induction of Foxp3 by TGF-B in conventional T cells^30^, Ikaros-deficient Treg exhibited normal Foxp3 expression. Instead, loss of Ikaros activity results in significant dysregulation of the Treg gene expression program, including pro-inflammatory cytokine and chemokine genes normally not expressed by Treg (*e.g*., *Ifng, Tnf, Il3, Il12rb, Tlr2*), and genes required for normal Treg function (*e.g*., *Tcf7, Lef1, Satb1, Nr4a1*). Many of these dysregulated genes are Foxp3 targets, and we show that Foxp3 cooperates with Ikaros to occupy the majority of Foxp3 binding sites in Treg.

Mice with Treg lineage-specific loss of Ikaros occupancy showed an accumulation of activated regulatory and helper T cells in the secondary lymphoid tissues, but no evidence for increased T cell infiltration into organs or frank autoimmunity. This might be explained by the fact that Treg isolated directly *ex vivo* showed elevated expression of many genes that promote Treg function, potentially balancing the dysregulated program predicted to have deleterious effects on Treg homeostasis and function. However, upon TCR stimulation, *Ikzf1*-deficient Treg induce a set of inflammatory Th1, Notch, and Wnt pathway genes normally repressed in Treg, and fail to control *in vivo* cellular and humoral immune responses mediated by conventional T cells. Importantly, our results show that the loss of suppressive function *in vivo* is not due to failure of Treg to home to sites of tissue inflammation. The lack of spontaneous inflammation or autoimmunity in *Ikzf1*-fl-Foxp3-Cre mice is similar to mice with Treg-specific deletion of *Prdm1*^31,32^, *Icos*^33^, *Il10*^34^ and *Mef2d*^35^. Mice lacking *Il10* in Treg exhibit mild colitis, but no autoimmunity, while mice with deletion of *Prdm1* in Treg show signs of autoimmunity only in aged mice. Moreover, deletion of *Mef2d, Blimp1, Icos,* or *Il10* in Tregs does not impact their suppressive activity *in vitro*, but impairs their function *in vivo* in the context of inflammation.

Although the literature is not in complete agreement, Eos and Helios are considered to be necessary for Treg function through their contribution to the Treg gene expression program. Eos, like Ikaros, is required to repress inflammatory gene expression by Treg^8,9^, and Eos can cooperate with Foxp3 to strengthen a core Treg gene expression program when ectopically co-expressed in conventional T cells^36^. Loss of Helios function in Treg has the primary effect of destabilizing *Foxp3* expression^7,11,12^, and also contributes to core Treg gene expression when ectopically co-expressed with Foxp3^36^. Despite these functions, neither Eos, Helios, nor Aiolos were able to compensate for the loss of Ikaros, despite the fact that *Ikzf1*-deficient Treg express all these proteins at comparable or higher levels as compared to wild-type Treg. The availability of Helios ChIP-seq^12^ and Helios-dependent gene expression^37^ data allowed us to compare Ikaros and Helios genome occupancy and gene regulatory programs (**Figure 10**). Of the 1838 Helios binding sites and 5477 Ikaros binding sites detected in Treg open chromatin, 64 are shared, representing only ∼3.5% of Helios binding sites and ∼1% of Ikaros binding sites. Similarly, we found a statistically significant overlap between the Ikaros-dependent set of 660 genes and the Helios-dependent set of 147 genes, but this consisted of only 9 genes (**Figure 10**). For five of these genes Ikaros and Helios have the same effect on expression, while Ikaros and Helios have the opposite effect on expression of the other four genes. This level of discordance at the level of both genome occupancy and gene regulation likely explains whey Helios, for example, cannot compensate for the loss of Ikaros in Treg.

**Figure 10.**
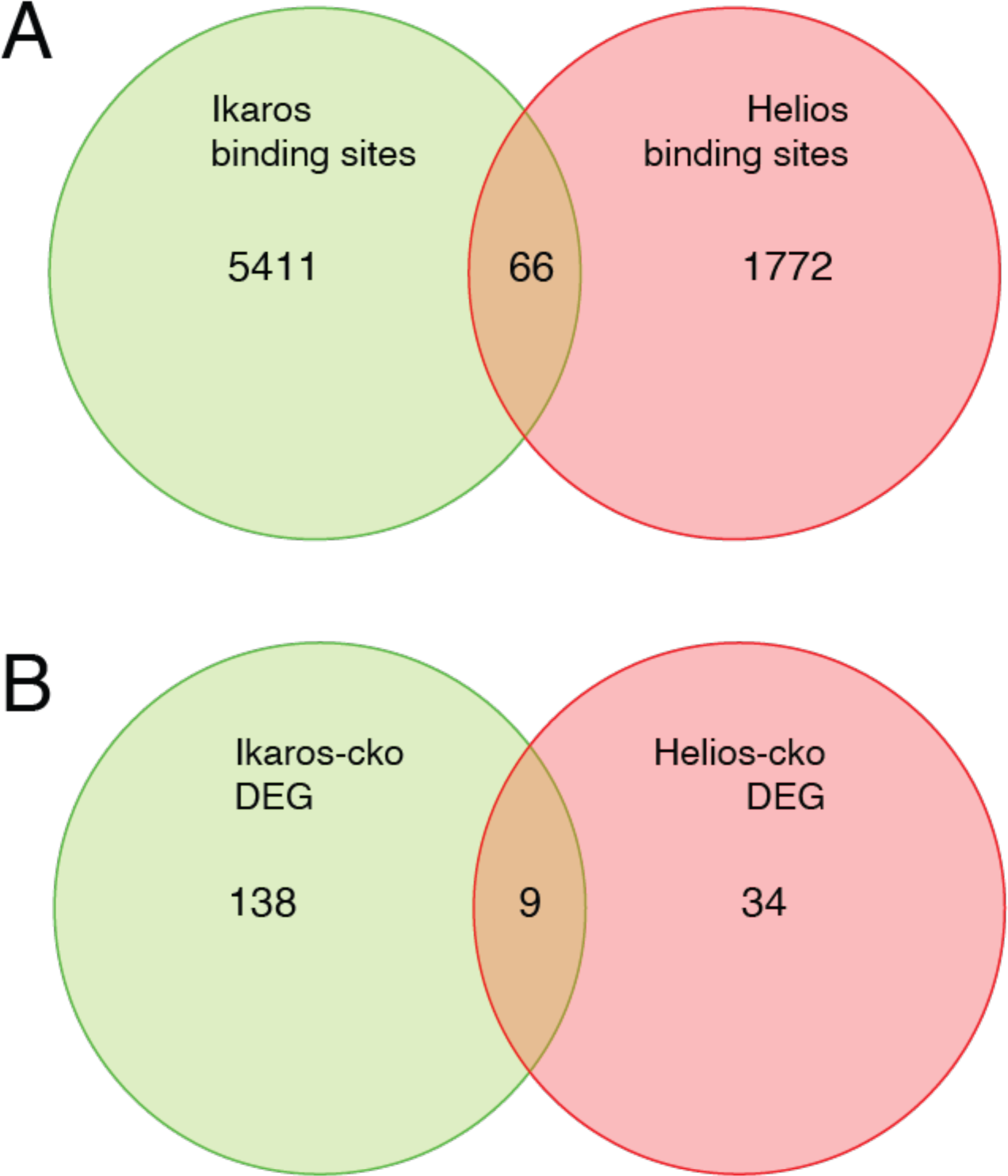
Comparison of Ikaros vs. Helios binding sites and regulated genes. Venn diagram in (**a**) depicts comparison of published ChIP-seq Helios binding sites^12^ (one replicate) with at least one bp of overlap with Ikaros binding sites from this study. Venn diagram in (**b**) depicts overlap of differentially expressed genes (FDR < 0.5 & abs(log2FC) > 1) in Ikaros or Helios conditional knockout^37^.

Coding mutations in *IKZF1* in humans is a cause of common variable immune deficiency (CVID) and autoimmunity^38^. To date, studies have focused on the impact of these mutations on T and B lymphocyte function, and our results here suggest Treg defects could also contribute to the immune dysregulation observed in these patients. In addition, common genetic polymorphism at the *IKZF1* locus have been associated with SLE susceptibility by GWAS^39^, and one mechanism for this is an SLE-associated distal regulatory element required for normal expression of Ikaros in human T cells^40^. Given the important roles for Ikaros in conventional B and T cell function, and its role newly defined here in regulatory T cell function, Ikaros is a relevant target for novel therapies for autoimmunity, organ transplant rejection, and cancer.

## Methods

### Antibodies

Fluorochrome-conjugated anti-mouse monoclonal antibodies CD3-AF700 (cat # 100216), CD4-BV785 (cat # 100453), CD8-PB (cat # 100725), CD25-BV650 (cat # 102038), GITR-PECy7 (cat # 120222), ICOS-PECy5 (cat # 107708), IFNg-PeCy7 (cat # 505826) and IL-2-PB (cat # 503820) were purchased from Biolegend. CD44-Percp-cyanine5.5 (cat# 45-0441-80, CD62L-APCeFL780 (cat # 47-0621-82), Foxp3-APC (cat# 17-5773-82), Eos-eFL660 (cat # 50-5758-80), Helios-PeCy7 (cat # 25-9883-42), Aiolos-PE (cat # 12-5789-80),and bCatenin-eFL660 (cat # 50-2567-42) were purchased from ThermoFisher Scientific. PD-1-PECy7 (cat# 25-9985-80), CXCR5-BV421 (cat # 562889), Bcl-6-PE (cat # 569522) and phosphor-STAT5-PE (pY694) (cat # 612567) were procured from BD Biosciences.

### Mice

The conditional *Ikzf1*-fl/fl mouse was provided by Dr. Meinrad Busslinger^13^. Mice were maintained at the Department of Veterinary Resource facility of CHOP. All animal experiments were performed according to protocols and guidelines approved by the CHOP animal care and use committee. To generate mice with conditional deletion of Ikaros in Tregs, homozygous *Ikzf1*-fl/fl mice were crossed with Foxp3-IRES-YFP-Cre mice^14^ purchased from JAX. Specific deletion of *Ikzf1* in Treg was confirmed by flow staining.

### Immunophenotyping and ELISA

Thymus, spleen and lymph nodes were collected from individual mice (5-7 weeks old) and single cell suspension was prepared in 1X PBS. RBC lysed lymphocytes were stained for CD3, CD4, CD8, CD25, CD44, CD62L, GITR, PD-1, ICOS, Foxp3, and Ikaros. For intracellular transcription factor staining, cells were fixed with eBioscience Perm/fix buffer (ThermoFisher Scientific). To determine Ikaros and Foxp3 expression, permeabilized cells were first incubated with rabbit anti-mouse Ikaros antibody (Abcam) diluted 1:2000 in perm/wash buffer for 1 hour and then cells were washed and stained with goat anti-rabbit-PE secondary antibody (1:2000 dilution) for 30 min. After washing, cells were stained for Foxp3, washed and analyzed by flow cytometry on a Cytoflex equipped for multicolor detection. Flow cytometry data analysis was conducted with Flowjo10 software. Secreted IL-2 and IFNg in cell culture supernatants were determined by ELISA following the instructions provided by the vendor, ThermoFisher Scientific.

### T cell and Treg purification

CD4+CD25- conventional T cells and CD4+CD25+ Tregs were purified from spleen and lymph node cell single cells suspension using Miltenyi Treg and CD4 purification kits. For FACS sorting, total CD4+ T cells were isolated first using Miltenyi CD4+ T cell purification kit and then sorted on a FACS Jaz sorter for CD4+ YFP+ Tregs.

### Cell culture

Purified Tregs were resuspended in RPMI 1640 medium supplemented with 10% FBS, 50 uM 2-ME, pencillin/ streptomycin and L-glutamine. Cells were stimulated with plate bound mouse anti-CD3 and anti-CD28 (1 ug/ml each) in 96 well plate and incubated at 37C in a cell culture incubator for the indicated times. For Treg proliferation assay, cells were labelled with Cell Trace (ThermoFisher Scientific) and stimulated with microbeads coated anti-mouse CD3/ CD28 (Dynabeads mouse cell activator). Cell proliferation was determined after 3 days of activation. PKC inhibitor, Calphostin C (PKF) was purchased from Cayman Chemical Company. Tregs were cultured in the presence of various concentrations of PKF in 96 well plate coated with anti-CD3 and anti-CD28 (1ug/ml each) for 3 days. Supernatant was collected for ELISA. For intracellular cytokine staining, cells were harvested and re-stimulated with PMA (15 ng), ionomycin (1uM), and Golgistop for 5 hrs. Cells were harvested, washed with 1X PBS and then stained for live cells with Live-dead aqua stain followed by staining for flow cytometry.

### STAT5 phosphorylation assay

Splenocytes were isolated from WT and Ikzf1-fl-Foxp3-YFP-Cre mice. Cells were washed with 1X PBS and pellet was resuspended in RPMI medium at 2×10^6^/mL. To induce STAT5 activation, aliquots of 10^6^ splenocytes were treated with recombinant mouse IL-2 (Sigma, cat # 11271164001) at 5-20 units/mL and cultured in 48 well plate for 30 minutes at 37C. Stimulated and unstimulated cells were harvested and washed with 2 mL of FACS buffer and then cells were fixed with BD transcription factor fixation/perm buffer (cat #562574, BD Bioscience) for 20 minutes at room temperature. Cells were washed with 2 mL of FACS buffer and cell pellet was fixed with 0.5 mL of 90% ice cold methanol for 30 min on ice. Cells were spun down, removed methanol and washed 2X with BD Perm/wash (1X) buffer. Cells were stained with an antibody cocktail prepared in 1X BD Perm/wash buffer containing fluorochrome-conjugated antibodies against CD4, CD44, CD62L, CD25, Foxp3, and phospho-STAT5 (pY694). Cells were stained at room temperature for 45 minutes followed by washing with BD Perm/wash buffer. Cell pellet was resuspended in 350 ul wash buffer and analyzed by flow cytometry. pSTAT5 staining was analyzed on gated Treg population.

### *In vitro* Treg suppression assay

Lymphocyte cell suspensions were prepared using the lymph nodes and spleen collected from the Foxp3-YFP-Cre, *Ikzf1*-fl-Foxp3-YFP-Cre, and C57BL/6 mice. Conventional CD4+ CD25-ve and CD4+ CD25+ Tregs cells were purified from the lymphocytes of wild-type and *Ikzf1*-cko mutant mice using Miltenyi Treg isolation kit (cat # 130-091-041). APCs were negatively selected from the lymphocytes of C57BL/6 mouse using Miltenyi CD90 (Thy1.2) cat #130-049-101 kit. APCs were gamma irradiated in a cesium irradiator. Ten million CD4 Tconv cells were labelled with CellTrace Violet and resuspended in RPMI medium at 1×10^6^ cells/mL. Labelled Tconv cells (50,000/well) were cocultured with 0.1×10^6^/well irradiated APCs plus various Treg:Tconv ratios (1:1, 1:2, 1:4, 1:8, 1:0) in 96 well round bottom plates. Cells were stimulated with soluble anti-CD3 (1ug/mL) and cells were cultured at 37C for 72 hours in a cell culture incubator. Cells were harvested, washed with 1X PBS and stained with live/dead aqua dye followed by flow staining with CD4, CD25, and CD44 flurochrome conjugated antibodies. Cells were analyzed on a Cytoflex flow cytometer and data was analyzed by Flowjo10 software. Cell division was quantified as described previously^50^, and percent suppression represents the reduction in cell division measured in the Tconv in the presence of Treg compared to no Treg.

### Co-immunoprecipitation analysis

293T cells were co-transfected with eukaryotic expression vectors encoding Flag-Ik1, Flag-Ik7, Foxp3, or control empty vector. A Flag-Runx1 construct was co-transfected with Foxp3 as a positive control for co-precipitation. After 48 hours of transfection, a cell lysate was prepared and Flag antibody immunoprecipitation was done for the lysates using a Flag IP kit (Zigma). Pulldown products were immunoblotted for Flag protein and Foxp3.

### T cell transduction

T cell transductions were performed as described previously^41^. Briefly, mouse CD4+ T cells were transduced with empty vector, Foxp3 vector or co-transduced with a retroviral vector expressing the dominant negative Ik7 isoform. After 3 days of transduction, cells were harvested and re-stimulated with plate bound anti-CD3 and anti-CD28. Supernatant was collected for IL-2 and IFNg ELISA and cells were harvested for Foxp3 chromatin immunoprecipitation.

### ChIP-seq library generation and analysis

For transcription factor and H3K27ac ChIP-seq analysis, we used *in vivo* expanded Tregs generated in mice using IL-2/anti-IL-2 complexes. Anti-mouse IL-2 antibody (BE0043) was purchased from Bioxcell and recombinant mouse IL-2 (carrier free, cat # 575408 from Biolegend). IL-2/anti-IL-2 complexes were prepared by mixing both reagents, incubating at 37C for 30 minutes, and diluted with 1X PBS. Foxp3-YFP-Cre and *Ikzf1*-fl-Foxp3-YFP-Cre mice (5-6 weeks old) were injected intraperitoneally with 200 ul of complex containing 2 ug IL-2 and 10 ug anti-IL-2. Each mouse received injection daily for 3 days and treatment-free for another 3 days before harvesting lymph nodes and spleens. For Ikaros ChIP-seq, total CD4+ T cells were negatively enriched using Miltenyi CD4 microbeads and then sorted for YFP+ Tregs by a FACSJaz sorter. Tregs purified through Miltenyi Treg purification kit were used for Foxp3, Ikaros, and H3K27ac-ChIP-seq. Naïve CD4+ T cells from Foxp3-YFP-Cre mice were also purified for Ikaros and H3K27ac ChIP-seq using a mouse CD4+ naïve purification kit purchased from Mitenyi Biotech. For all ChIP-seq experiments, 3 biological replicates of cells isolated from 3 individual mice were used. Chromatin immunoprecipitations were performed using ChIP-IT high-sensitivity kits (cat #53040, Active Motif) following the manufacturer’s instructions. Briefly, 5×10^6^ purified cells were fixed in medium for 15 minutes at room temperature using complete cell fixative solution prepared using formaldehyde and cell fixative solution from the kit. The reaction was stopped by adding 1/20 media volume of stop solution, cells were washed with ice-cold PBS and a cell pellet was stored at -80C for later use or cells were lysed in chromatin preparation buffer cells as described in the protocol. The cell pellet was resuspended in ChIP buffer and chromatin was sonicated by a QSonica Q800R sonicator with settings: amplitude 20%, pulse for 30 seconds on, 30 second off, for a total of 30 cycles. For input DNA preparation, 25 ul of the sonicated sample was removed and DNA was isolated as suggested in the protocol. An agarose (1%) gel electrophoresis was performed for DNA isolated from the input fraction to determine the sonication efficiency. ChIP-validated anti-mouse Ikaros antibody (cat #39355) and H3K27Ac antibody (cat #39133) were purchased from Active Motif. For Foxp3 ChIP-seq, eBioscience anti-mouse monoclonal antibody (cat#14-5773-82) was purchased from ThermoFisher Scientific. The volume of the sheared chromatin was adjusted to 200 ul using ChIP buffer, and to which was added 5 ul of Protease inhibitor, and a mix containing ChIP antibody (4 ug) and 5 ul of blocker, mixed and pre-incubated at room temperature for a minute. Final volume of the ChIP reaction was 240 ul, which was incubated at 4C overnight on a rotator. Antibody precipitated chromatin immune complexes were collected using washed protein G agarose beads and immune complexes were washed 5X in ChIP filtration columns using wash buffer, and eluted the DNA with elution buffer. The eluted DNA was reverse cross linked and further purified through DNA purification columns. ChIP’d DNA was eluted from the column using 30 ul of DNA purification elution buffer. All ChIP-seq and input DNA libraries were made using ThruPLEX DNA-Seq kit (cat #R400674, Takara Bio, USA) following the manufacturer’s instructions. In brief, the fragmented DNA obtained from ChIP reaction or input DNA was end repaired to generate blunt ends, to which stem-loop adaptors with blocked 5’ ends are ligated. Libraries were amplified through high fidelity amplification buffer mix and Takara dual indexing primers (cat# R400407). Finally, the amplified dual indexed libraries were purified using AMpure XP beads (Beckman Coulter, Cat # A63880) at 1:1 ratio. The purified DNA was recovered from the beads using 20 ul of TE buffer. The library quality was checked on a bioanalyzer using high sensitivity DNA Chip. Library DNA concentration was determined using Qubit. Dual-indexed ChIP-seq libraries were pooled and sequenced on a the Illumina NovaSeq 6000 platform. Reads were aligned to mm9 using bowtie2 and duplicated reads were marked using Picard with parameters VALIDATION_STRINGENCY=LENIENT and ASSUME_SORTED=true and removed samtools. Library quality was accessed using samtools flagstat to assess library complexity and strand cross correlation to assess^42^. Peaks were called using MACS2 with the parameters -g mm9 –nomodel –p 0.01 –keep-dup_all with the – extsize estimated fragment size from the strand cross correlation for each replicate of H3K27ac, Ikaros, or Foxp3 with matching input sample. Reads were subsequently filtered by the ENCODE mm9 blacklist regions. Within condition peaks were filtered to ones found in at least two replicates. Binary comparison between different ChIP peaks and ATAC-seq was performed using the R package GenomicRanges (1.46.1) findOverlaps function. For differential comparisons of H3K27ac ChIP signal, reads were normalized against background (10K bins) using csaw (v1.28, http://bioconductor.org/packages/release/bioc/html/csaw.html). Peaks with a cpm value less than 3.0 were removed from further differential analysis. Differential analysis was performed using glmQLFit approach in edgeR (v3.36.0) with the normalization scaling factors calculated from csaw. FDR < 0.05 was used as the cutoff for statistical significance. Signal reproducibility between replicate samples was accessed using pairwise pearson correlation tests and principal component analysis. Significant OCR overlapping with H3K27ac peaks were annotated as enhancers. Correlations between enhancer accessibility, H3K27ac ChIP signal, and expression of nearest gene were computed using pearson correlation coefficient implemented in the R function cor.test. Super-enhancers were called using the rank ordering of super-enhancer (ROSE) algorithm^43^. Briefly OCR called by ATAC-seq were used as input regions and clustered by genomic coordinates with a 12.5kb stitching window. Merged replicates of WT and Ikzf1-cko H3K27ac signal and input were used as measure of enhancer activity. The signal is represented as input subtracted reads per million per basepair and then are ranked-ordered. The position in the ranked list where the change in signal (slope when x-axis is the super-enhancer rank and y-axis is signal) equals 1 is used to define super-enhancers by the rapid increase in enhancer activity. Super-enhancers were defined independently for WT and Ikzf1-cko H3K27ac data. ChIP peaks were annotated to their nearest gene based on linear genomic distance.

### RNA-seq library generation and analysis

Total CD4+ T cells isolated from individual WT YFP Cre+ and Ikzf1-Treg-cko mice (3 biological replicates) were sorted for YFP+ Tregs on a FACS Jaz sorter. For stimulation, Tregs were activated with plate bound anti-CD3 and anti-CD28 (1 ug/ml each) for 4 hrs. Total RNA was isolated from the unstimulated and stimulated Tregs using Direct-zol RNA micro prep kit (Zymo Research). Quality of the DNase treated total RNA was checked on a bioanalyzer. Ribosomal RNA was depleted from the total RNA using QIAseq fast select multi-RNA removal kit for mouse RNA(Qiagen) and then RNAseq libraries were made using NEB Next Ultra II Directional RNA library prep kit for Illumina. RNAseq library quality was checked on a high sensitivity bioanalyzer and dual indexed libraries were sequenced to 51 bp reads on the Illumina NovaSeq 6000 platform. The pair-end fastq files were mapped to genome assembly mm9 by STAR (v2.6.0c)^44^ for each replicate. Ensembl v67 mm9 annotation was used for gene feature annotation and the unadjusted read count for gene feature was calculated by htseq-count (v0.6.1)^45^ with parameter settings -f bam -r pos -s reverse -t exon -m union. The gene features annotated as rRNAs were removed from the final sample-by-gene read count matrix. The differential analysis was performed in R (v3.3.2) using the edgeR package (v3.16.5)^46^. Briefly, the raw reads on genes features with total CPM (read counts per million) value of less than 3.66 (the bottom 25% gene features when comparing the highest count per condition across all samples) were removed from differential analysis. The trimmed mean of M-values (TMM) method were used to calculate normalization scaling factors and quasi-likelihood negative binomial generalized log-linear (glmQLFit) approach was applied to the count data and through pairwise comparisons of stimulated and unstimulated IK cko and WT. The differential expression genes (DEGs) between were identified with cut-off FDR < 0.05 and absolute logFC>1. TPM values were calculated for differentially expressed genes and scaled expression values (across rows) were depicted using the R package ComplexHeatmap (2.10.0). Immunologic signature gene sets annotated in MSigDB (v7.0) were used for gene set enrichment analyses. Statistical significance of gene set enrichment for up and down regulated were determined using the hypergeometric test (one-tailed), implemented in the R phyper function.

### ATAC-seq library generation and analysis

YFP+ Tregs were FACS sorted from Foxp3-YFP-Cre and *Ikzf1*-fl-Foxp3-YFP-Cre mice. Half were stimulated with plate bound anti-CD3+ anti-CD28 for 4 hours and half were left unstimulated. One hundred thousand cells were lysed with 50 ul of cold lysis buffer (10mM Tris pH 7.4, 10 mM NaCl, 3mM MgCl2, 0.1% IGEPAL CA-630) and centrifuged at 550g for 10 minutes at 4C. Supernatants were discarded and nuclear pellets were subjected to Tn5 transposition using Nextera DNA preparation kits (cat #1502812). DNA was purified from transposition reaction using Qiagen Min-Elute PCR purification kits. Transposed DNA fragments were PCR amplified and indexed using Nextera index kits. PCR reaction products were size selected using AMpure-XP beads and DNA fragments were re-suspended in 10 mM Tris-HCl. DNA concentration was determined by Qubit, and ATAC seq library quality was checked on a bioanalyzer. Dual indexed libraries were sequenced on the Illumina NovaSeq 6000 platform. Open chromatin peaks were called using the ENCODE ATAC-seq pipeline (https://www.encodeproject.org/atac-seq/). Briefly, pair-end reads from three biological replicates for each cell type were aligned to hg19 genome using bowtie2, and duplicate reads were removed from the alignment. Narrow peaks were called independently for each replicate using macs2 (-p 0.01 --nomodel --shift -75 --extsize 150 -B --SPMR --keep-dup all --call-summits) and ENCODE blacklist regions (ENCSR636HFF) were removed from peaks in individual replicates. Peaks from all replicates were merged by bedtools (v2.25.0) within each cell type and the merged peaks present in less than two biological replicates were removed from further analysis. Finally, ATAC-seq peaks from both cell types were merged to obtain reference open chromatin regions. Quantitative comparisons of wild-type and *Ikzf1*-cko open chromatin landscapes were performed by evaluating read count differences against the reference OCR set. De-duplicated read counts for OCR were calculated for each library and normalized against background (10K bins of genome) using the R package csaw (v1.8.1). OCR peaks with less than 3.6 CPM support in the top 25% of samples were removed from further differential analysis. Differential analysis was performed independently using edgeR (v3.16.5). Differential OCR between cell types were called if FDR<0.05 and absolute log2 fold change >1.

### Transcription factor motif enrichment

Enrichment of known transcription factors binding motifs was determined for the differential sets of OCRs using the R package PWMEnrich (v4.30.0). Enrichment of differential OCRs were calculated using all OCRs in the background model. Sequences were extracted from the bioconductor genome reference BSgenome.Mmusculus.UCSC.mm9.masked (v1.3.99) using the R package Biostrings (v2.62.0). We used JASPAR2020 position-weight matrix database as the motif reference^47^. P values were adjusted with using FDR. Transcription factor footing improves the confidence of TF binding over pure sequencing matching. We identified putative Ikaros and Foxp3 footprint using HINT-ATAC (<u>H</u>mm-based Ide<u>N</u>tification of <u>T</u>ranscription factor footprints). HINT-ATAC corrects for Tn5 cleavage bias using a HMM based approach to identify de novo TF footprints. Replicate deduplicated ATAC-seq bam files were merged and used as input to identify TF footprints located in the consensus set of OCRs. The de novo TF footprints were then matched to known TF PWMs in the JASPAR2020 database.

### DNA methylation analysis

To assess the natural thymic or peripheral origin of Tregs, Foxp3 TSDR region which is fully demethylated in natural Tregs, was analyzed for CpG methylation by sodium bisulfite sequencing method. DNA methylation level at IFNg intronic enhancer was also analyzed. Briefly, 1 ug of DNA extracted from FACS sorted CD4+ YFP+ Tregs or CD4 conventional T cells was bisulfite converted following^48^. The converted DNA was desalted using the Wizard DNA clean-up system (Promega), desulphonated, neutralized and precipitated with ethanol. The Foxp3 CNS2-TSDR, the IFNg promoter, and the IFNg intronic enhancer regions were PCR amplified from bisulfite converted DNA using nested PCR primers as described^15,49^. Gel purified PCR products were cloned into PGEM-T easy vector, and plasmid DNA individual clones were sequenced using SP6 primers.

### Adoptive transfer colitis model

To assess the *in vivo* suppressive capacity of Ikaros deficient Tregs, adoptive T cell transfer experiments were set up with Rag1 KO mice. CD4+ CD25-ve conventional T cells and Tregs were purified from the splenic and lymph node lymphocytes of male WT YFPCre+ or IKZF1 FL/FL Foxp3 Cre+ mice. Batches of Rag1 KO male mice consisting of 5 mice per group were received retro-orbital injection of 1 million WT CD4 conv T cells along with (0.25X 10^6) WT or Ikaros deficient Tregs. Recipient mice were weighed 3 times a week and observed for various IBD induced clinical symptoms. Mice were sacrificed after 40 days of T cell transfer, spleen, mesenteric lymph nodes and colon were removed. Small pieces of tissue were cut from the lower part of colon and were fixed in formal fixative for histopathological analysis. Single cell suspension was made from spleen and mesenteric lymph nodes from individual mice, lymphocyte cell density was estimated using a hemocytometer. Lymphocytes were stained for CD4, CD8, CD25, CD44 and Foxp3 and analyzed by flow cytometry. Absolute T cell count was estimated using hemocytometer count and cell frequency derived from the flow cytometry analysis. Variation in weight curve between groups were statistically analyzed by ANOVA (GraphPad Prism).

### Cardiac transplantation

Fully MHC-mismatched BALB/c hearts (H-2d) were transplanted heterotopically into Foxp3-YFP-Cre and *Ikzf1*-fl-Foxp3-YFP-Cre mice by anastomosis of the donor ascending aorta and pulmonary artery to the recipient infrarenal aorta and pulmonary artery. On days 0, 2 and 4 of posttransplant, the transplant recipient mice were given i.p injection of CTLA4 Ig fusion protein (200 ug) and CD154 (200 ug). A separate batch of cardiac recipient mice were administered donor specific transfusion (DST) with Balb/c splenocytes (5 million cells/recipient) and a single dose of MR1 (250 ug). Cardiac graft survival was determined by abdominal palpation, and cessation of cardiac contraction was considered as rejection of the graft. Grafts were monitored for a 100 days, and graft survival data were analyzed by Kaplan-Meir/log-rank methods. For histopathological analysis, a separate batch of mice consisting of 3 mice per group were subjected to same transplant procedure for a short duration and then these recipient mice were sacrificed, grafts were removed before rejection, fixed in 1X formal solution, and analyzed by standard histopathological methods.

### Cardiac histopathology and intragraft gene expression analysis

For histopathologic analysis of cardiac graft and colon from IBD experiments, portions of tissues were fixed in Shandon formal-Fixx solution (1X), ThermoScientific. Tissues were processed and embedded in paraffin. Histologic sections were cut at 4um thickness and stained with hematoxylin and eosin (H&E) stain and or with alcian blue. Immunohistochemical staining was performed for CD4 and Foxp3 by the CHOP Pathology cores. Blinded histopathological evaluations were performed by a histopathologist. For intragraft gene expression analysis and histopathological analysis, a separate batch of mice consisting of 3 mice per group were subjected to same transplant procedure with a duration of 19 days for CTLA4Ig and MRI treated mice and a duration of 14 days for MRI and DST treated mice, based on the weakness of transplant heart palpitation. These transplant recipient mice were sacrificed, graft was removed, a portion of the cardiac graft was frozen immediately in liquid nitrogen and a second portion of the graft tissue was fixed in 1X formal solution and analyzed by standard histopathological methods. Total RNA was extracted from the homogenized graft tissue using Trizol. One microgram of RNA was treated with DNase to avoid DNA contamination. It was then reverse transcribed using iScript cDNA synthesis kit (cat #170-8891) purchased from Bio-Rad. Gene expression analysis for the cardiac graft was performed by qRT PCR using Fast SYBR Green mastermix on a Applied Biosystems step one plus real time PCR system. Following primer pairs were used for the qRT PCR amplification: Foxp3-Exp-F; AAAAGGAGAAGCTGGGAGCTATG, Foxp3-Exp-R; GTGGCTACGATGCAGCAAGAG, IFNg-Exp-F; TTGCCAAGTTTGAGGTCAACAA, IFNg-Exp-R; GCTGGATTCCGGCAACAG, TNF-a-F; CTGTAGCCCACGTCGTAGC, TNF-a-R; TTGAGATCCATGCCGTTG, IL-6-F; TGTTCTCTGGGAAATCGTGGA, IL-6-R; CTGCAAGTGCATCATCGTTGT, CXCR3-F; TACCTTGAGGTTAGTGAACGTCA, CXCR3-R; CGCTCTCGTTTTCCCCATAATC, CXCL-10-F; TGCCGTCATTTTCTGCCTCA, CXCL-10-R; GGACCGTCCTTGCGAGAG, Bcl-6-F; GTGTCCCCCAGTTTGTGTCA, Bcl-6-R; TGGAGCATTCCGAGCAGAAG, DGKA-F; CAAACAGGGCCTGAGCTGTA, DGKA-R; CGAGACTTGGCATAGGTGCT, CD27-F; GGATGTGTGAGCCAGGTACA, CD27-R; GGGTGTGGTAGTCTGGAGAG, m18S RNA-F; TTCGAACGTCTGCCCTATCAA, m18S RNA-R; ACCCGTGGTCACCATGGTA.

## Acknowledgments

Author contributions: Conceptualization: ADW, RMT, MCP; Methodology: RMT, LW, MCP; Investigation: ADW, RMT, LW, MCP; Visualization: RMT, MCP, WWH; Supervision: ADW, WWH, SFAG; Writing—original draft: RMT, MCP; Writing—review & editing: ADW, WWH. Funding provided by the Fred & Susanne Biesecker Pediatric Liver Center and the Center for Spatial and Functional Genomics at The Children’s Hospital of Philadelphia. The authors have no competing interests to declare. All data needed to evaluate the conclusions in the paper are present in the paper and/or the Supplementary Materials. Genome-scale datasets are available on the Gene Expression Omnibus, accession GSE200179.

---

**Supplementary Figure 1.**
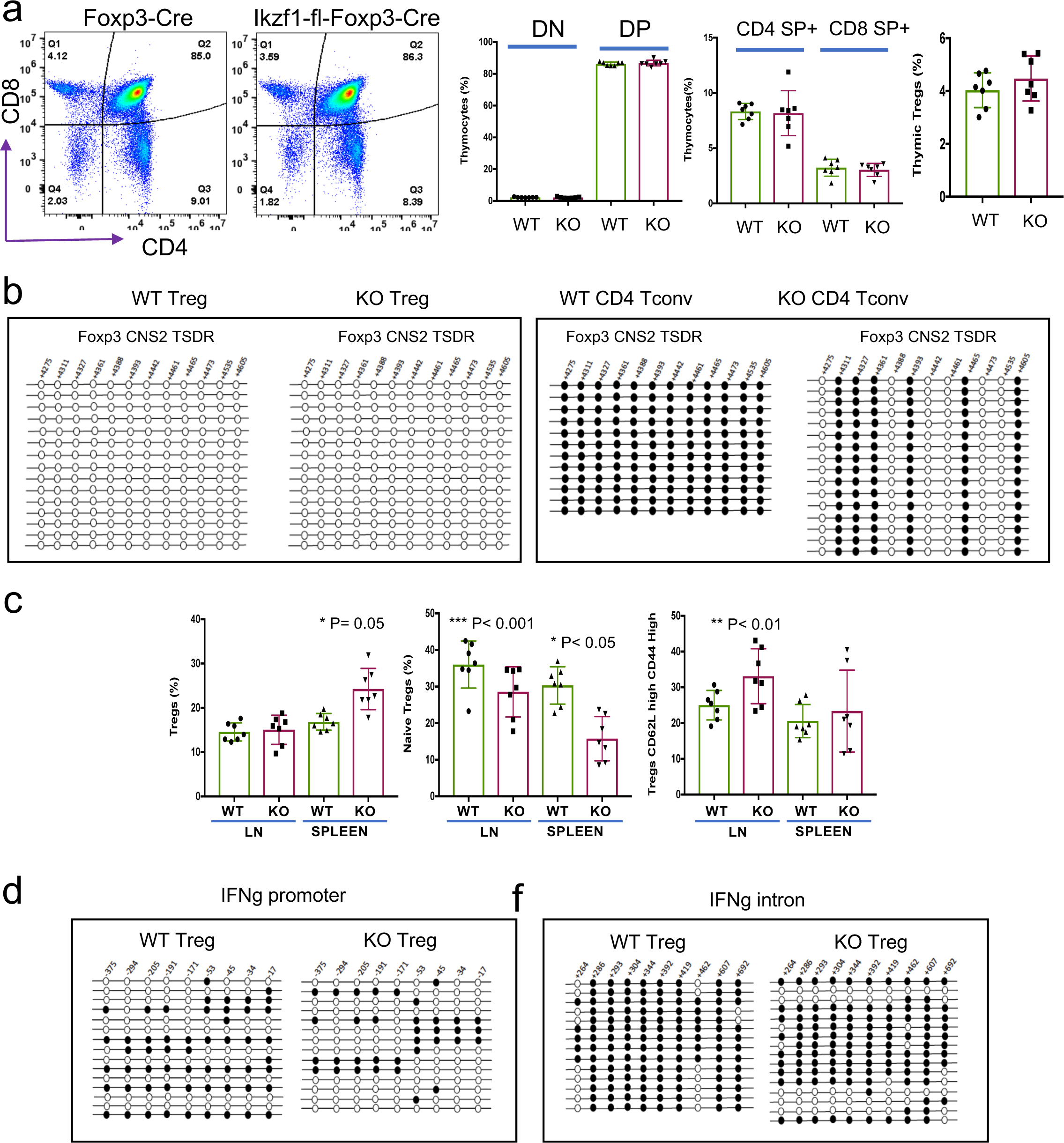
Immunophenotyping of thymi from *Ikzf1*-fl-Foxp3-YFP-Cre or Foxp3-YFP-Cre control mice (**a**). Bisulfite DNA methylation analysis of the Foxp3 CNS2/TSDR region in peripheral Treg and Tconv from *Ikzf1*-fl-Foxp3-YFP-Cre or Foxp3-YFP-Cre control mice (**b**). Quantification of total peripheral Treg, CD44^lo^ cTreg, CD44^hi^ eTreg, and PD1^hi^CXCR5^hi^ Tfreg in *Ikzf1*-fl-Foxp3-YFP-Cre and Foxp3-YFP-Cre mice (**c**). Bisulfite DNA methylation analysis of the IFNg promoter and IFNg intronic enhancer (**d**) regions in peripheral Treg and Tconv from *Ikzf1*-fl-Foxp3-YFP-Cre or Foxp3-YFP-Cre control mice.

**Supplementary Figure 2.**
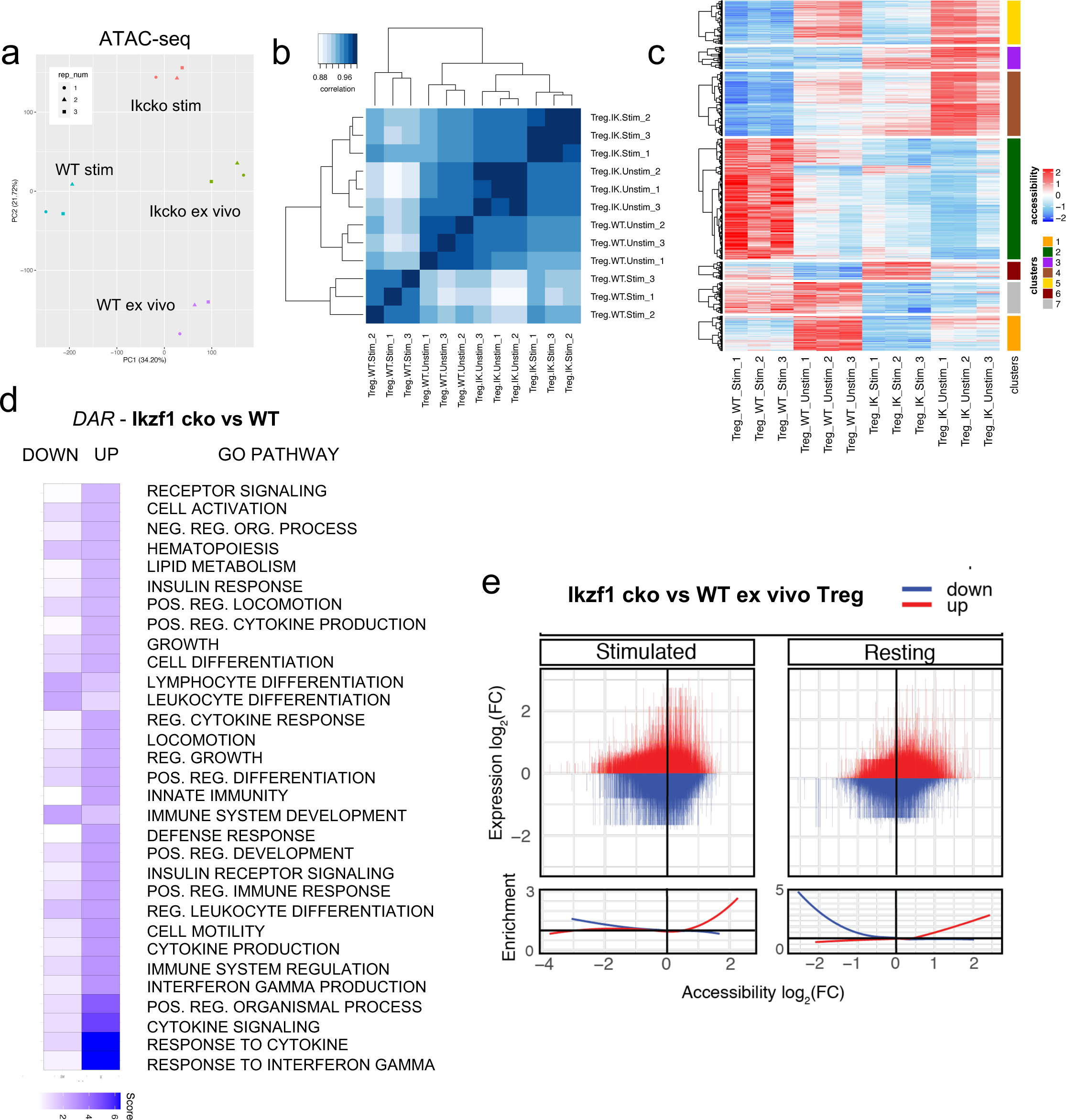
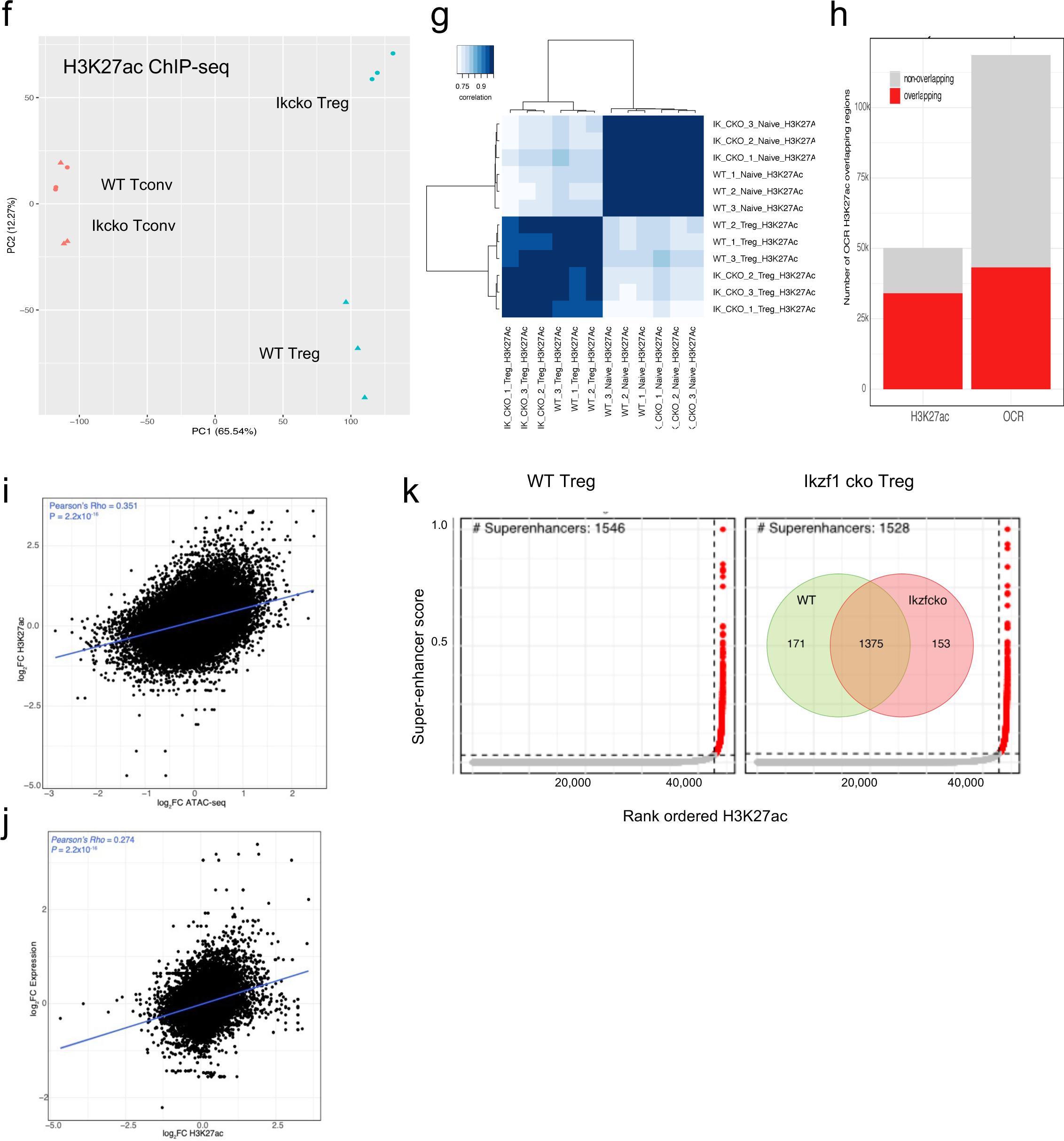
ATAC-seq and H3K27ac ChIP-seq library metrics. Assessment of WT and *Ikzf1*-cko *ex vivo* and stimulated Treg library reproducibility by principal component (**a**) and spearman correlation (**b**) analysis. (**c**) Hierarchical clustering of 12,030 differentially accessible regions (DAR, FDR<0.05, abs[logFC]>1) into 7 groups from the root. Red vs. blue indicate increased vs. decreased accessibility. (**d**) Enrichment of GO terms for genes nearest to DAR (FDR<0.05, >5 genes). (**e**) Association of DAR accessibility changes with changes in expression of the nearest gene. Enrichment is over the mean increase or decrease in expression, indicated by red vs. blue. Assessment of WT and *Ikzf1*-cko H3K27ac ChIP-seq library reproducibility by principal component (**f**) and pairwise spearman correlation (**g**) analysis. Overlap (**h**) and Pearson correlation between ATAC-seq and H3K27ac ChIP-seq peaks (**i**) and and H3K27ac and gene expression (**j**). Identification of super-enhancer regions based on H3K27ac density (**k**). The Venn diagram (inset) depicts the number of super-enhancers shared between WT and *Ikzf1*-cko Treg (orange), unique to WT Treg (green), and unique to *Ikzf1*-cko Treg (red).

**Supplementary Figure 3.**
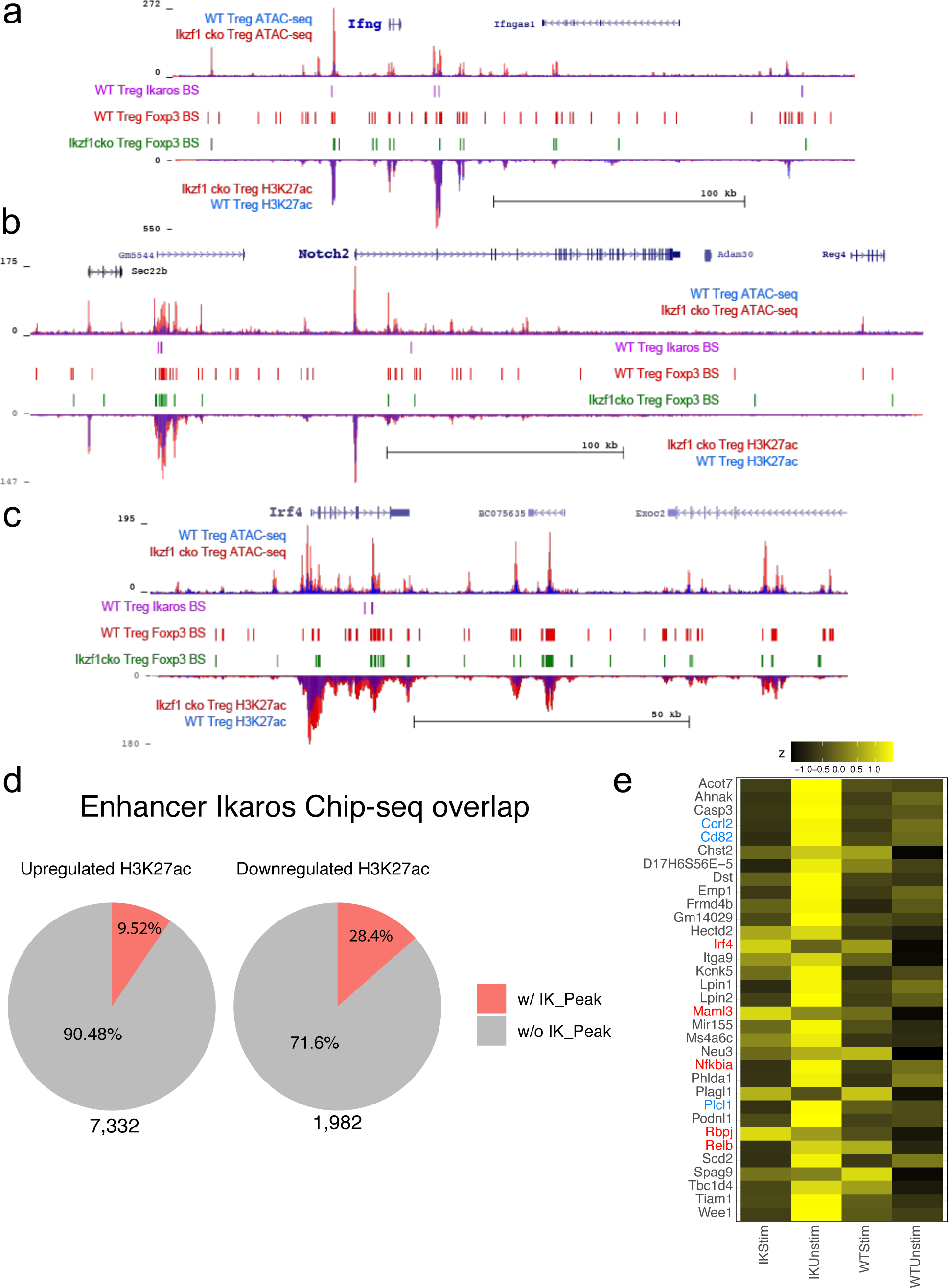
Multi-omic integration of differential ATAC-seq, ChIP-seq and RNA-seq analyses. (**a-c**) Ikaros-dependent changes in accessibility, enhancer signatures, and Foxp3 occupancy at genes differentially expressed in *Ikzf1*-cko Treg. Open chromatin (top tracks) and H3K27ac (bottom tracks) in WT (blue) and *Ikzf1*-cko (red) Treg, Ikaros binding sites (purple marks) and Foxp3 binding sites (red marks) in WT Treg, and Foxp3 binding sites in *Ikzf1*-cko Treg (green marks) at the *Ifng* (**a**) and *Notch2* (**b**) and *Irf4* (**c**) loci. (**d**) Overlap between Ikaros ChIP-seq peaks and differentially acetylated regions is indicated in red in the pie charts. (**e**) Differential H3K27ac (z-score) at top Ikaros-bound DEG in ex vivo and stimulated WT and *Ikzf1*-cko Treg.

**Supplementary Figure 4.**
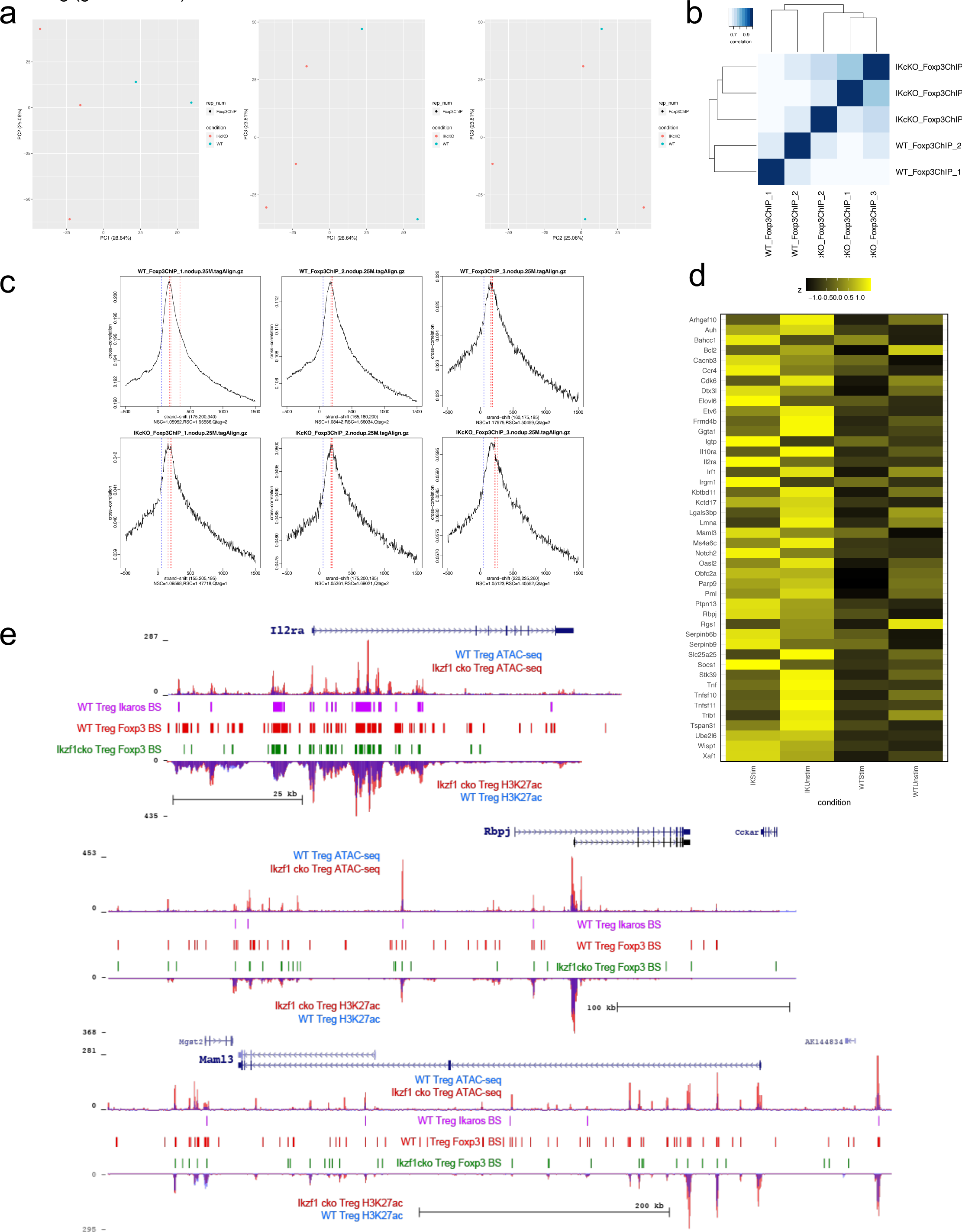
Foxp3 ChIP-seq library metrics. Assessment of WT and *Ikzf1*-cko library reproducibility by principal component (**a**) and pairwise spearman correlation (**b**) analysis. (**c**) Strand cross-correlation analysis of FoxP3 ChIP-seq signal vs. noise corresponding to the broad peak (red) and phantom peak (blue; mappability bias). (**d**) Differential H3K27ac (z-score) at top Foxp3-bound DEG in ex vivo and stimulated WT and *Ikzf1*-cko Treg. (**e**) Ikaros-dependent changes in accessibility, enhancer signatures, and Foxp3 occupancy at the *Il2ra, Rbpj,* and *Maml3* genes differentially expressed in *Ikzf1*-cko Treg. Open chromatin (top tracks) and H3K27ac (bottom tracks) in WT (blue) and *Ikzf1*-cko (red) Treg, Ikaros binding sites (purple marks) and Foxp3 binding sites (red marks) in WT Treg, and Foxp3 binding sites in *Ikzf1*-cko Treg (green marks).

**Supplementary Figure 5.**
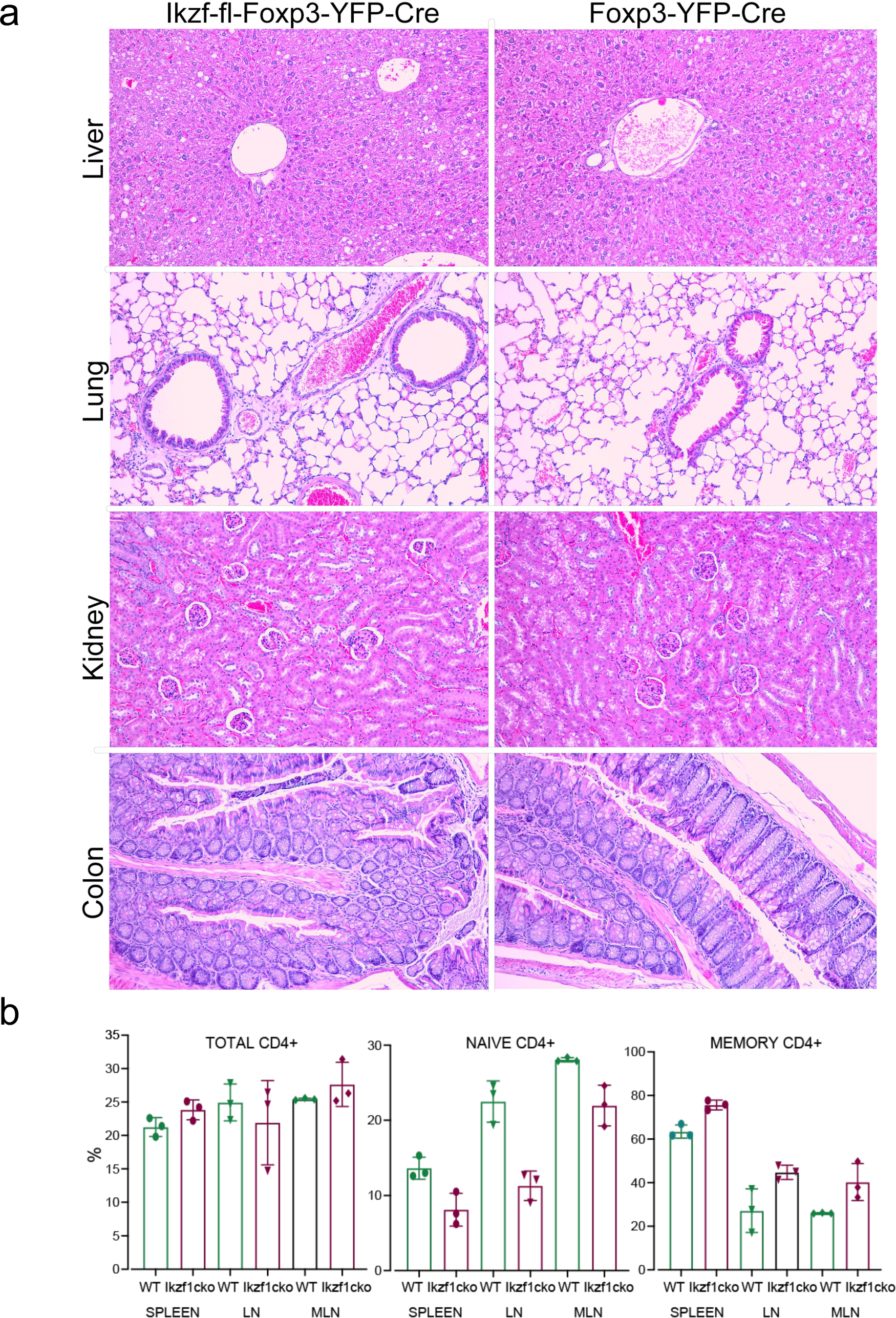
(**a**) Analysis of H&E-stained paraffin sections of liver, lung, kidney, and colon from 1 year-old *Ikzf1*-fl-Foxp3-YFP-Cre (N=3) or age-matched Foxp3-YFP-Cre control mice (N=3). Representative images at 100X magnification are shown. All tissues showed normal appearance with no evidence of spontaneous autoimmunity. (**b**) Immunophenotyping of Tconv from *Ikzf1*-fl-Foxp3-YFP-Cre and Foxp3-YFP-Cre mice. Frequencies of total (left panel), CD62L^hi^CD44^lo^ naive (middle panel), and CD62L^lo^CD44^hi^ memory (right panel) phenotype Tconv in secondary lymphoid tissues of 10 month-old *Ikzf1*-fl-Foxp3-YFP-Cre (purple) and Foxp3-YFP-Cre (green) mice (N=3).

**Supplementary Figure 6.**
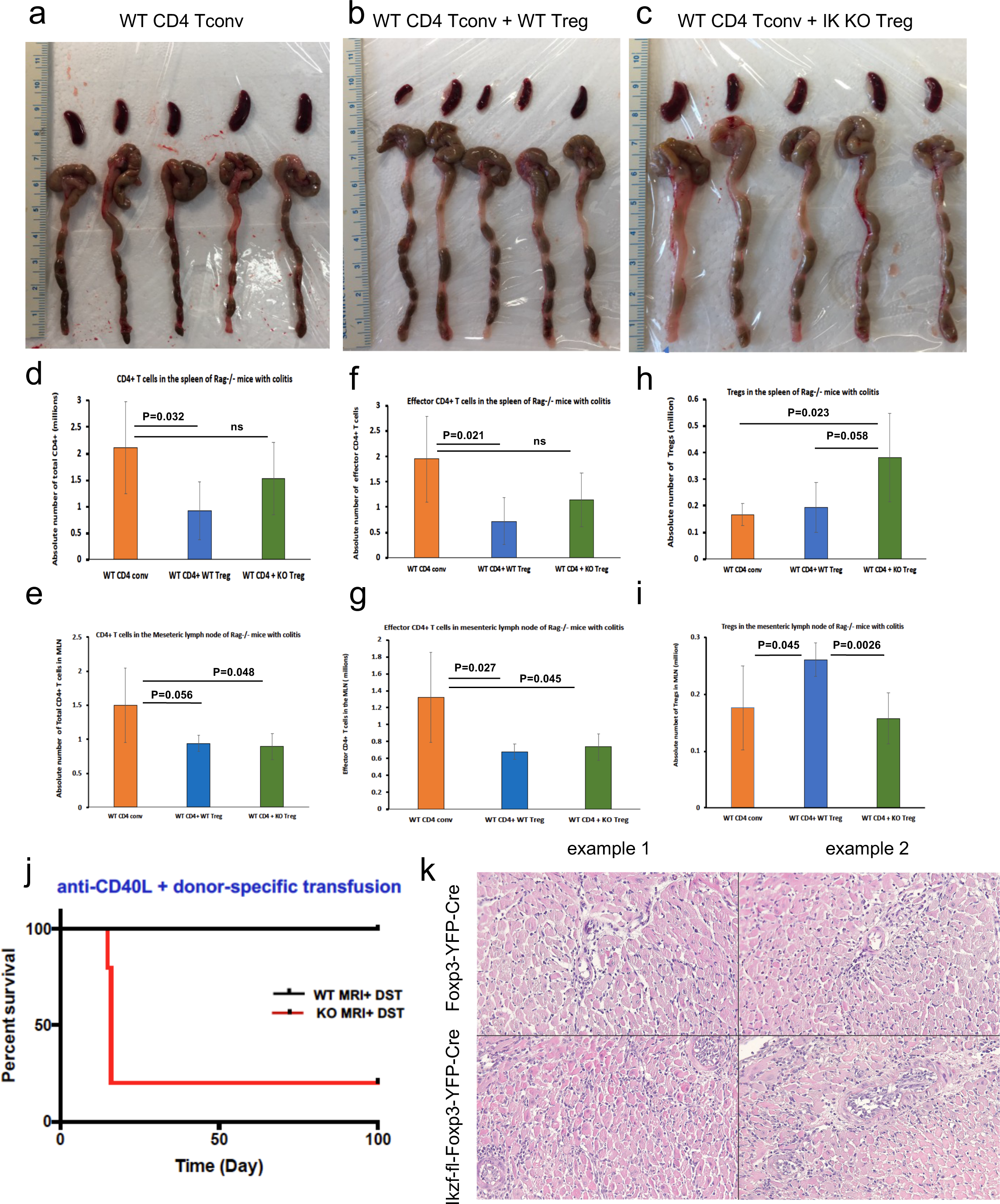
Role of Ikaros in Treg-mediated protection from colitis. Spleens and colons from Rag1ko recipients of WT Tconv (**a**), WT Tconv plus WT Treg (**b**) and WT Tconv plus *Ikzf1*-cko Treg (**c**) harvested at day 40 post-transfer. Quantitation of total CD4+ T cells (**d,e**), CD4+CD44+ effector Tconv (**f,g**), and CD4+Foxp3+ Treg (**h,i**) in the spleens (**d,f,h**) and mesenteric lymph nodes (**e,h,i**) from Rag1ko recipients of WT Tconv (orange), WT Tconv plus WT Treg (blue) and WT Tconv plus *Ikzf1*-cko Treg (green) harvested at day 40 post-transfer. Role of Ikaros in Treg-dependent acquired cardiac transplant tolerance. (**j**) B6 *Ikzf1*-fl-Foxp3-YFP-Cre (red) or Foxp3-YFP-Cre (black) mice received BALB/c cardiac allografts, donor-specific transfusion (DST), and anti-CD40L. Graft survival was monitored for 100 days. (**k**) Histopathological analysis of cardiac grafts harvested at day 14 post-transplant from Foxp3-YFP-Cre and *Ikzf1*-fl-Foxp3-YFP-Cre recipients (N=3, scale=200x).

## Supplementary Table files

Table S1: Differential analysis of RNA-seq data

Table S2: Differentially expressed genes with known roles in Treg function

Table S3: Gene ontology of differentially expressed genes

Table S4: Differentially accessible regions

Table S5: Differential analysis of H3K27 acetylation

Table S6: Super-enhancer analysis

Table S7: ChIP-seq analysis of Ikaros binding

Table S8: Transcription factor motif enrichment

Table S9: ChIP-seq analysis of Foxp3 binding

